# Ocean-wide comparisons of mesopelagic planktonic community structures

**DOI:** 10.1101/2021.02.26.433055

**Authors:** Janaina Rigonato, Marko Budinich, Alejandro A. Murillo, Manoela C. Brandão, Juan J. Pierella Karlusich, Yawouvi Dodji Soviadan, Ann C. Gregory, Hisashi Endo, Florian Kokoszka, Dean Vik, Nicolas Henry, Paul Frémont, Karine Labadie, Ahmed A. Zayed, Céline Dimier, Marc Picheral, Sarah Searson, Julie Poulain, Stefanie Kandels, Stéphane Pesant, Eric Karsenti, The Tara Oceans coordinators, Peer Bork, Chris Bowler, Colomban de Vargas, Damien Eveillard, Marion Gehlen, Daniele Iudicone, Fabien Lombard, Hiroyuki Ogata, Lars Stemmann, Matthew B. Sullivan, Shinichi Sunagawa, Patrick Wincker, Samuel Chaffron, Olivier Jaillon

## Abstract

For decades, marine plankton have been investigated for their capacity to modulate biogeochemical cycles and provide fishery resources. Between the sunlit (epipelagic) layer and the deep dark waters, lies a vast and heterogeneous part of the ocean: the mesopelagic zone. How plankton composition is shaped by environment has been well-explored in the epipelagic but much less in the mesopelagic ocean. Here, we conducted comparative analyses of trans-kingdom community assemblages thriving in the mesopelagic oxygen minimum zone (OMZ), mesopelagic oxic, and their epipelagic counterparts. We identified nine distinct types of intermediate water masses that correlate with variation in mesopelagic community composition. Furthermore, oxygen, NO_3_ ^-^ and particle flux together appeared as the main drivers governing these communities. Novel taxonomic signatures emerged from OMZ while a global co-occurrence network analysis showed that about 70% of the abundance of mesopelagic plankton groups is organized into three community modules. One module gathers prokaryotes, pico-eukaryotes and Nucleo-Cytoplasmic Large DNA Viruses (NCLDV) from oxic regions, and the two other modules are enriched in OMZ prokaryotes and OMZ pico-eukaryotes, respectively. We hypothesize that OMZ conditions led to a diversification of ecological niches, and thus communities, due to selective pressure from limited resources. Our study further clarifies the interplay between environmental factors in the mesopelagic oxic and OMZ, and the compositional features of communities.

## Introduction

Below the ocean’s sunlit layer lies the mesopelagic zone that occupies around 20% of the global ocean volume [1]. The mesopelagic zone is biologically defined as starting where photosynthesis no longer occurs (<1% irradiance; around 200m depth), down to its lower boundary where there is no detectable sunlight (around 1000m depth) [2]. This twilight ecosystem cannot rely on photoautotrophy, but sustains its energetic requirements by the combination of heterotrophic, chemoautotrophic, and chemo-mixotrophic metabolisms, together with physicochemical processes. Among the latter, the fraction of upper ocean productivity that escapes epipelagic recycling and sinks by gravity or is delivered by the daily vertical migration of zooplankton constitutes an essential energy source in deep waters and is a vector for attached organisms [3].

Considerable attention has been devoted to the mesopelagic layer in recent years, given its recognized potential for exploitation for bioresources and fisheries [4], potentially becoming an important source of goods for the global bioeconomy [5]. So far, efforts have been made to increase the knowledge of mesopelagic macrofauna by studying the abundance and diversity of nekton. Concerning the mesopelagic community’s microscopic fraction, previous reports have shown a stratification of planktonic communities by the water column. In this regard, the mesopelagic zone displays a distinct assemblage of dsDNA viruses [6], Nucleo-Cytoplasmic Large DNA viruses [7], prokaryotes [8–10], and eukaryotes [11]. These studies have highlighted a global organization different from that of the surface. For example, the mesopelagic plankton diversity does not show a latitudinal diversity gradient trend from pole-to-pole, peaking at lower latitudes [12], and also displays a higher heterogeneity compared to epipelagic waters [13]. Conversely, the mesopelagic microbiome seems to make crucial links in the food web between phototrophic primary production from the sunlit layer and dark ocean specialized consumers [10,14,15].

Among the studies conducted in mesopelagic zones, particular efforts have been made to explore regions characterized by extreme conditions, such as oxygen minimum zones (OMZs). These zones are formed by relatively old, slowly upwelling waters, often lying below highly productive surface zones [16], and are currently increasing in volume in the ocean [17]. OMZ prokaryotic communities are well documented and taxa such as *Nitrospira*, *Marinimicrobia*, and anammox bacteria from the phylum Planctomycetes have been reported as typical taxonomic features for OMZ regions studied so far [18–23]. In contrast, knowledge about viruses and eukaryotic diversity in OMZs is still rudimentary. A prevalence of specific eukaryotic taxa such as Ciliophora, Dinoflagellata, MALV, and Acantharia has been reported, together with a higher metabolic activity of these taxa [24–26]. Viruses may have a key role in OMZ ecosystem feedback by modulating the local community (host-virus relationship) [27–29].

The last decades have seen a significant increase in large-scale oceanic surveys [30–32]. Despite the advances reported in previous studies [10,33,34], most mesopelagic community studies have been limited to geographically or ecologically fragmented regions, or to specific taxonomic groups, mainly because of the inherent difficulties of accessing this zone on a global scale [35]. Moreover, the combination of biotic and abiotic factors influencing community structure, [36,37], has been poorly explored in the mesopelagic zone. Extending this knowledge by including more comprehensive and homogeneous datasets from assorted geographical and oceanographic systems should lead to a global understanding of this layer. Understanding plankton community structure and dynamics is fundamental to anticipate the impact of global warming and acidification in these regions.

Here, we present a trans-kingdom omics-based comparative study of epipelagic, oxic- and OMZ-mesopelagic communities. To this end, we compiled *Tara* Oceans survey taxonomic DNA barcodes data from four oceanographic basins using standardized sampling protocols. In particular, we focused on mesopelagic environmental drivers, ecology, and taxa associations networks in both oxic and OMZ.

## Materials and Methods

### Sample collection and pre-processing

The environmental and biological data were obtained during the *Tara* Oceans expedition (2009-2012) in 32 oceanographic stations located in the Indian Ocean (IO - 037, 038, 039), Pacific Ocean (PO - 097, 098, 100, 102, 106, 109, 110, 111, 112, 122, 131, 132, 133, 135, 137, 138), South Atlantic Ocean (SAO - 068, 070, 072, 076, 078) and North Atlantic Ocean (NAO - 142, 143, 144, 145, 146, 148, 149, 152) comprising tropical and subtropical regions (Figure 1). Physico-chemical environmental data were obtained along a vertical profile at each station. Temperature, salinity, and oxygen were measured using a CTD-rosette system with a coupled dissolved oxygen sensor. Chlorophyll-*a* concentrations were measured using high-performance liquid chromatography. Nutrient concentrations were determined using segmented flow analysis. All these metadata are available at PANGAEA [38–43] (https://doi.pangaea.de/10.1594/PANGAEA.875582).

**Figure 1:**
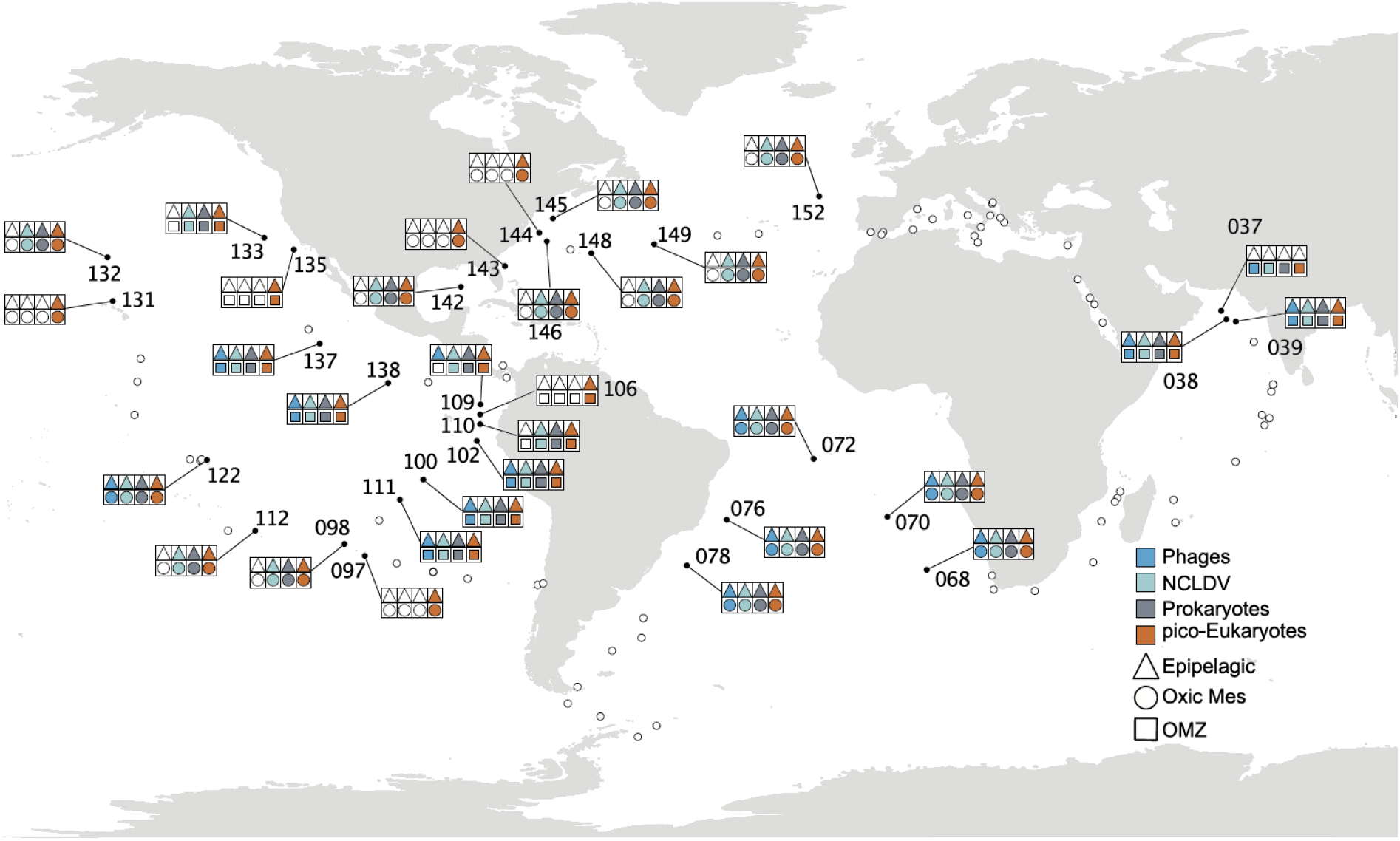
Geographical locations of *Tara* Oceans epipelagic and mesopelagic sampling sites included in this study. Symbol colors represent organism groups evaluated in the present study.

The vertical distribution of marine particles was investigated with an Underwater Vision Profiler (UVP, [44,45]) mounted on the CTD-Rosette. The UVP acquires images in a coherent water volume (1 L) delimited by a light sheet issued from red light-emitting diodes. Automatic identification of objects was made using Ecotaxa [46], based on a learning set of visually identified, manually classified objects and associated features. Images were classified to distinguish mesozooplankton from non-living objects and artifacts (e.g., detrital particles, fibers, and out-of-focus objects).

Water vertical profiles of temperature and salinity generated from the CTD were used to identify the water masses by plotting a temperature x salinity (T/S) diagram using the Ocean Data View V 5.0 (ODV) software package [47].

Three different water layers were sampled: surface (SRF, 3-7 m), deep chlorophyll maximum (DCM - depth identified according to the peak of chlorophyll-*a* fluorescence obtained *in situ*), and mesopelagic (ranging from 200-1000 m) [48]. The planktonic community was sampled by partitioning the pumped seawater by filtering each sampled depth with different filter sizes, the on-board sampling methodology is detailed by Pesant et al. [42].

Among the sampled mesopelagic zones, 13 of them were identified as deficient in oxygen and classified as oxygen minimum zone (OMZ, stations IO - 037, 038, 039 / PO - 100, 102, 106, 109, 110, 111, 133, 135, 137, 138). The OMZ were categorized as suboxic: <10 μM O_2_/kg seawater and anoxic: (<0.003 μM/kg seawater or undetectable with most sensitive techniques, e.g., STOX sensors) [Units of O_2_ concentration: 1 mL.L^-1^=1.43 mg. L^-1^ or 1 mL.L^-1^=44.64 μM] [25].

Our dataset comprises different organismal size-fractions from viruses (two dsDNA-virus bacteriophage families *Podoviridae* and *Myoviridae* - hereafter named as phages and Nucleo-Cytoplasmic Large DNA viruses - hereafter named as NCLDV) to pico-eukaryotes.

Phage libraries were constructed from seawater samples filtered at 0.22 μm, concentrated using iron chloride flocculation, and treated with deoxyribonuclease (DNase). NCLDV *polB* and prokaryotic 16S rRNA gene sequences were extracted from plankton metagenomes sequenced from 0.22–1.6 or 0.22-3 μm filters, and the pico-eukaryote dataset was obtained by V9-18S rRNA gene marker amplification from 0.8-3 or 0.8-5 μm filters. Details of sample preparation and sequencing procedures are fully described in Alberti et al. [49].

Phage read counts was accessed through the search for the marker genes *gp23* (*Myoviridae*) and *polA* (*Podoviridae*) in the protein collection GOV2.0 derived from metagenomic sequencing described in Gregory et al. [6]. The NCLDV read counts profile was obtained from the *polB* marker gene gathered from the OM-RGC.v2 catalog [9] as described in Endo et al. [7]. The Prokaryotic read counts was assessed from a metagenomic dataset called 16S mitag, as described in Sunagawa et al. [8], which does not rely on PCR amplification [50]. Sequences matching “Eukaryota”, “chloroplasts”, and “mitochondria” were removed from the final Prokaryotic OTU table. Clustering and annotation of pico-eukaryote V9-18S rRNA gene PCR-amplicons are described in de Vargas et al. [51], and functional annotation of taxonomically assigned V9-18S rRNA gene metabarcodes was improved afterwards; in this case, we conserved in the final data only sequences assigned to the “Eukaryota” domain. More details concerning the acquisition and pre-processing of the sequence data used in the present study were compiled from earlier publications and are provided in Supplementary Methods. Throughout the manuscript, we use the classical term “OTU” to denote taxonomic units, although we did not employ the clustering technique historically associated with this acronym. More comments about “OTU” usage are available in Supplementary Methods.

To simplify the biological information and obtain a concise epipelagic dataset (EPI), we merged redundant SRF and DCM OTUs by summing their read counts for each taxonomic group and preserved non-redundant OTUs. Putative biases produced by this merging procedure were tested and discarded as shown in the Supplementary Methods; they did not affect the conclusions.

Afterward, to deal with the compositional nature of the data, OTU count matrices (EPI and MESO) were transformed using robust CLR (centered log ratio) after adding the value of one as a pseudo count. Robust CLR transformation considers only values greater than 0 to calculate the geometric mean and avoid biases due to sparse data [52].

### Epipelagic and Mesopelagic Community and Environmental differences

We applied an NMDS analysis based on the Bray-Curtis dissimilarity matrix on CLR transformed data. The ‘metaNMDS’ function from the vegan R package [53] was applied to confirm community differences between epipelagic and mesopelagic layers. Homogeneity of the sampled environmental parameters was checked using the ‘betadisper’ function (homogeneity of multivariate dispersions in the vegan package). The analysis was conducted using the Euclidean distance matrix of the environmental variables with the depths (epipelagic, mesopelagic) as group factor. A permutation test statistically confirmed the results.

### Ecological inferences and statistics

Ecological patterns were inferred using environmental variables to constrain the variation observed in biological data (CLR transformed) for planktonic samples using Canonical Correspondence Analysis (CCA) in the vegan R package. A set of physico-chemical variables for the discrete depths were selected for the ecological inferences, such as nitrate (NO_3_ ^-^), oxygen, temperature, salinity, density, and particles using particle flux UVP data. In order to avoid collinearity among factors, the selected variables were checked for variance inflation factor using the vif.cca function and tested for significance by ANOVA-like tests performed by ‘anova’ implemented in vegan with 999 permutations. The significance of the effect of each variable was tested individually using all others parameters as covariables (independently from their order in the model) by applying the option ‘margin’ to the ‘anova’ function in vegan.

Permutational multivariate analysis of variance (PERMANOVA) was performed with the function ‘adonis’ in vegan to determine the relationship between mesopelagic community composition and predefined water masses based on 999 permutations.

### Classification of organisms Eco-region

In order to detect organisms specific to epipelagic (EPI), oxic mesopelagic (Oxic MES), and OMZ eco-regions, only sampling sites containing both epipelagic and mesopelagic information were considered: in total, 25 stations for NCLDV, prokaryotes, and pico-eukaryotes, and 13 for phages. We ran a Kruskal-Wallis test (‘kruskal.test’ from stats R package [54]) to detect differential OTU abundances between eco-regions, followed by a Benjamini & Hochberg correction to avoid false discovery rate (FDR - p.value.bh). Organisms with a p-value <0.05, indicating a difference within groups, were subject to a post-hoc Dunn test (‘dunn.test’ from dunn.test R package [55]) to identify preferential eco-regions for each OTU. From these results, OTUs non-significant Kruskal-Wallis tests (p.value.bh >0.05) were assigned to the “ubiquitous” group. In contrast, those with significant p-values.bh were classified as EPI, Oxic MES, or OMZ if only the corresponding eco-region was elected according to the Dunn test. Organisms with no significant differences between Oxic MES and OMZ were assigned to Core MES.

### Co-occurrence network inference

For investigation of ecological associations between organisms across eco-regions, a co-occurrence network was inferred. In this analysis, phage samples were not included due to the lower number of stations sampled. Therefore, samples for NCLDV, prokaryotes and pico-eukaryotes from stations 038, 039, 068, 070, 072, 076, 078, 098, 100, 102, 109, 110, 111, 112, 122, 132, 133, 137, 138, 142, 145, 146, 148, 149 and 152 were retained. OTUs with a relative abundance lower than10^-4^ and counting fewer than 5 observations were discarded. Network inferences were performed on CLR transformed data using FlashWeave version 0.18 implemented in Julia version 1.2 [56], using the sensitive and heterogeneous mode. FlashWeave assumes features to be multivariate Gaussian distributed in CLR-transformed space.

We analyzed this global co-occurrence network by delineating communities (or modules) using the Clauset-Newman-Moore algorithm [57]. These modules are subsets of OTUs, obtained by maximizing the co-occurrences within modules and minimizing connections between them. Next, we investigated modules enriched in OTUs from specific eco-regions using Fisher’s exact test using the “fisher.test” function from the stats R package, followed by the Benjamini & Hochberg correction to control the FDR due to multiple testing.

## Results and Discussion

Leveraging the resources produced by the *Tara* Oceans project, we deciphered differences between epipelagic and mesopelagic beta-diversity stratification, with a particular emphasis on the role of environmental variables such as temperature, oxygen, salinity, NO_3_ ^-^, chlorophyll-*a*, and particle flux (see Methods). Here, we combined the diversity information obtained for surface and DCM samples to consider a single epipelagic group. We observed a clear distinction between the mesopelagic and epipelagic communities without loss of signal, as shown in previous studies [6–11], supporting our subsequent analyses reported here (Supplementary Figure 1, Supplementary Methods Figures 1 and 2). Consequently, we first investigated differences among physicochemical characteristics of the mesopelagic and epipelagic sampling sites. We observed a high dissimilarity gradient among sites for both layers (Supplementary Figure 2a, b). Mesopelagic samples were heterogeneously distributed, with most of the points placed distant from the group centroid (located in the center of the cloud of points identified for each group) (Supplementary Figure 2a). In contrast, epipelagic points displayed a large variance due to the samples positioned apart from the main cluster (Supplementary Figure 2a). These results underlie the heterogeneity of environmental conditions encountered in both sampled layers, and this environmental variation may be an important factor that can directly influence community composition.

**Figure 2:**
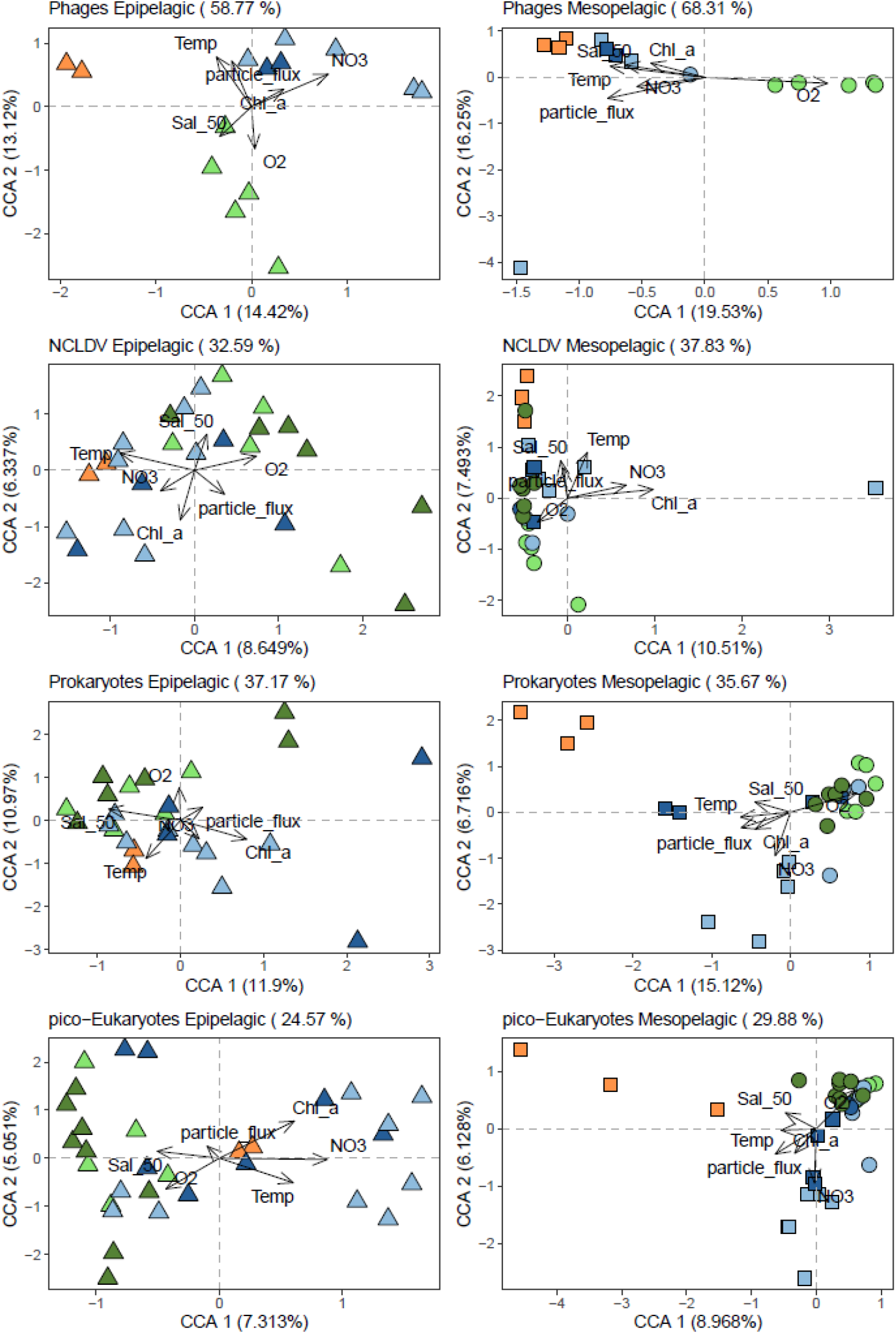
Ordination plot of canonical correspondence analysis (CCA) from epipelagic (left) and mesopelagic (right) communities based on OTU composition. Percentages in parentheses are the amount of variation constrained - in titles represent the total in each analysis, and in the axis represent the correspondent value for each dimension. Arrows represent environmental quantitative explanatory variables with arrowheads indicating their direction of increase. Shapes represent sampling sites. Shape formats represent eco-regions, epi: epipelagic, Oxic MES: oxic mesopelagic, OMZ: oxygen minimum zone mesopelagic. IO: Indian Ocean, NAO: North Atlantic Ocean, NPO: North Pacific Ocean, SAO: South Atlantic Ocean, SPO: South Pacific Ocean.

Next, to quantify how much of the differences in the assemblages’ variance can be explained by environmental conditions, we employed canonical correspondence analysis (CCA) using the environmental variables measured at discrete depths as constraint variables. The results showed that abiotic factors explained 40.5% on average of community variance for both layers (Figure 2). The phage assemblage was the exception, for which about 58% of the epipelagic variation and 68% of the mesopelagic variation could be explained by the variables investigated (Figure 2). Our analysis also demonstrated a clustering according to the different oceanic basins studied for all the assemblages, confirmed by a PERMANOVA analysis (Figure 2, Supplementary Table S1). A basin-scale biogeographical structure was already shown for virus, bacteria and protist in the epipelagic layer [58,59]. Here, we showed that this structuration appears even more pronounced in the mesopelagic samples at global scale.

Furthermore, we assessed the variance of communities regarding each environmental parameter as explanatory variables individually. In contrast with epipelagic communities mainly structured by temperature, as observed elsewhere [6,8,11,12,60,61], temperature was only a significant variable structuring the viruses (phage and NCLDV) in the mesopelagic layer. However, oxygen, NO_3-_ and particle flux appeared as common environmental drivers governing at least three out of four mesopelagic assemblages (Table 1, complete analysis in Supplementary Material Table S2).

**Table 1.** ANOVA p-value of the variance explained by environmental variables for global model including all tested environmental variables (Global) and for each explanatory variable individually with all the others used as covariables (independently from their order in the model)

| Assemblages | Depth | Global | Temp °C | Salinity | O<sub>2</sub> [μmol/Kg] | NO<sub>3</sub><sup>-</sup> [μmol/L] | Chl-a [μmol/m<sup>3</sup>] | Particle flux |
| --- | --- | --- | --- | --- | --- | --- | --- | --- |
| Phages | Epi | 0.001 | 0.048 | 0.132 | 0.126 | 0.001 | 0.308 | 0.013 |
| Phages | MES | 0.001 | 0.006 | 0.151 | 0.001 | 0.230 | 0.306 | 0.001 |
| NCDLV | Epi | 0.001 | 0.001 | 0.033 | 0.036 | 0.083 | 0.019 | 0.197 |
| NCDLV | MES | 0.002 | 0.043 | 0.186 | 0.005 | 0.031 | 0.001 | 0.048 |
| Prokaryotes | Epi | 0.035 | 0.062 | 0.088 | 0.303 | 0.152 | 0.106 | 0.606 |
| Prokaryotes | MES | 0.006 | 0.410 | 0.590 | 0.004 | 0.039 | 0.575 | 0.418 |
| pico-Eukaryotes | Epi | 0.025 | 0.108 | 0.576 | 0.373 | 0.065 | 0.173 | 0.807 |
| pico-Eukaryotes | MES | 0.001 | 0.168 | 0.304 | 0.033 | 0.001 | 0.016 | 0.009 |

Previous studies have identified oxygen as one of the main drivers of the eukaryotic community structure in OMZ regions [25,26,62]. These studies mainly compared community composition along the oxygen gradient within the water column depth, from the surface downwards. However, depth stratification of plankton communities is evident even in regions with high oxygen concentrations, so distinct parameters co-varying with depth must be taken into account in addition to the oxygen gradients [63].

In addition to the physicochemical parameters, our results show that particle flux (derived from UVP measurements) was a significant variable structuring phage, NCLDV and pico-eukaryote mesopelagic assemblages (Table 1, complete analysis in Supplementary Material Table S2). These observations support previous reports about the high correlation of this environmental factor with phages, finding possible relevance for the carbon pump’s functioning in epipelagic layers [14]. This observation may also reflect the association with virus (phages and NCLDV) inputs from overlaying water layers via sinking particles [64,65]. Furthermore, Bettarel et al. [66] suggested that marine aggregates can act as virus-factories, where these entities use the adsorbed bacteria to replicate and therefore be massively exported through the water column (one-way motion). The authors also showed that adsorbed bacteria can easily detach from aggregates (two-way motion), which can explain the lack of correlation between prokaryotes and particle flux observed here. On the other hand, pico-eukaryotes were also driven by particle flux. Durkin et al. [67], demonstrated that about 25% of epipelagic diversity can be detected on marine sinking particles and that the particle associated diversity is linked to the size and type of particle (fecal pellet loose or dense, aggregates and detritus).

*In situ* physico-chemical measurements have revealed the dynamics and fluctuating nature of the ocean, even over a short time scale [68]. The heterogeneity in mesopelagic layers given by deep currents, the impact of surface production, and the low mixing levels may favor a diversification in the mesopelagic community living in different water masses, leading to species adaptation-acclimation. The *Tara* Oceans expedition route included samples from common or distinct water masses defined by temperature/salinity profiles - T/S, comprising regionally connected or unconnected stations. We identified nine different water masses in the mesopelagic sampled locations (Figure 3). We could confirm significant differences among mesopelagic communities sampled in these different water masses based on the PERMANOVA test (Table 2). This result indicates that the oceanic patchiness created by distinct water masses significantly shape community beta-diversity in the mesopelagic layer, which would imply it to be a critical component for mesopelagic community variation for all the assemblages studied (phages, NCLDV, prokaryotes, and pico-eukaryotes). Thus, we hypothesize that this result may be explained by two non-exclusive causes related to water masses: (i) past common origin among water masses that have drifted or (ii) constant connectivity by ocean circulation between sampled sites belonging to the same water mass.

**Figure 3:**
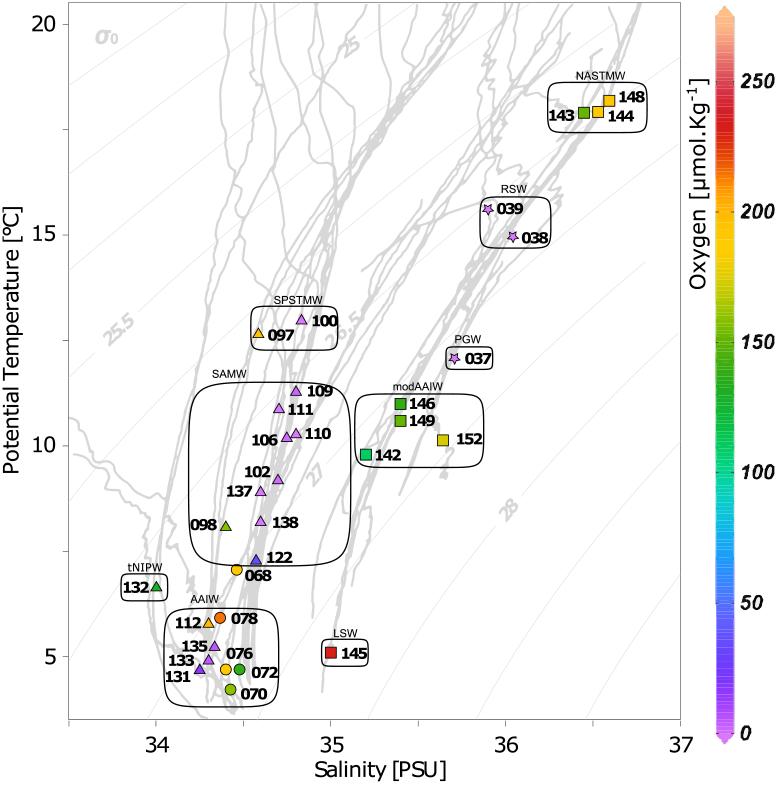
Temperature and salinity plot indicating water mass designation for all mesopelagic samples. Formats represent the different oceanic basins (▪ - North Atlantic Ocean, ● - South Atlantic Ocean, ▴- Pacific Ocean, ★- Indian Ocean). Colors indicate the oxygen concentration at the sampling depth. LSW - Labrador Sea Water; AAIW - Antarctic Intermediate Water; tNPIW transitional North Pacific Intermediate Water; SAMW - Subantarctic Mode Water; SPSTMW - South Pacific Subtropical Mode Water; modAAIW - modified Antarctic Intermediate Water; PGW - Persian Gulf Water mass; RSW - Red Sea Water mass; NASTMW - North Atlantic Subtropical Mode Water.

**Table 2.**
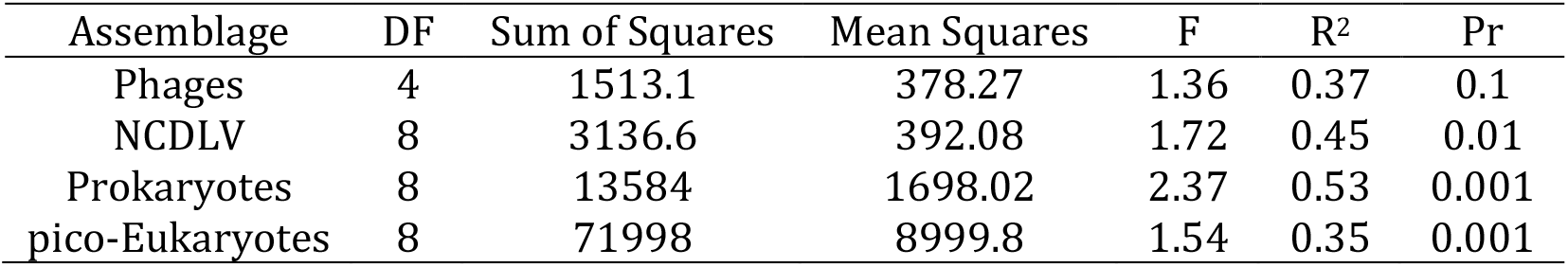
Proportion of the assemblages variation explained by water masses using the Permutation multivariate analysis of variance (PERMANOVA)

| Assemblage | DF | Sum of Squares | Mean Squares | F | R<sup>2</sup> | Pr |
| --- | --- | --- | --- | --- | --- | --- |
| Phages | 4 | 1513.1 | 378.27 | 1.36 | 0.37 | 0.1 |
| NCDLV | 8 | 3136.6 | 392.08 | 1.72 | 0.45 | 0.01 |
| Prokaryotes | 8 | 13584 | 1698.02 | 2.37 | 0.53 | 0.001 |
| pico-Eukaryotes | 8 | 71998 | 8999.8 | 1.54 | 0.35 | 0.001 |

We addressed another lingering question, resolving planktonic community signatures of oxic MES and OMZ regions from those observed in epipelagic layer. For this, we classified OTUs into three eco-regions: 1) EPI, 2) Oxic MES, and 3) OMZ. OTUs were classified as Core MES when commonly present in Oxic MES and OMZ samples. Taxa that were either equally abundant in all three eco-regions or not statistically confirmed to a single eco-region were classified as ubiquitous (Supplementary Figure 3, Supplementary Tables S3-S7). Using this approach, we could identify ubiquitous taxa that are likely to thrive in a wide range of environmental conditions, or that may be detected in mesopelagic samples due to the simple vertical movement of sinking particles. This classification should help avoid putative biases inherent to the metabarcoding methodology.

More specifically, we were able to identify Oxic MES and OMZ signatures mainly at the infra-taxonomic level (OTU-species) for all biotic groups investigated (Figure 4, Supplementary Figures 3-7, Supplementary Tables S3-S7). This reflected the wide ecological niche occupied by the different species at a higher taxonomic level (i.e. family). At the species level, we observed large taxonomic plasticity of OTUs that occurred equally in both Oxic MES and OMZ samples, called Core MES. However, most OTUs are not yet classified at the infra-taxonomic level (Supplementary Tables S3-S7). This observation reflects the knowledge gap about the biodiversity and functional plasticity of species thriving in this ecosystem.

**Figure 4:**
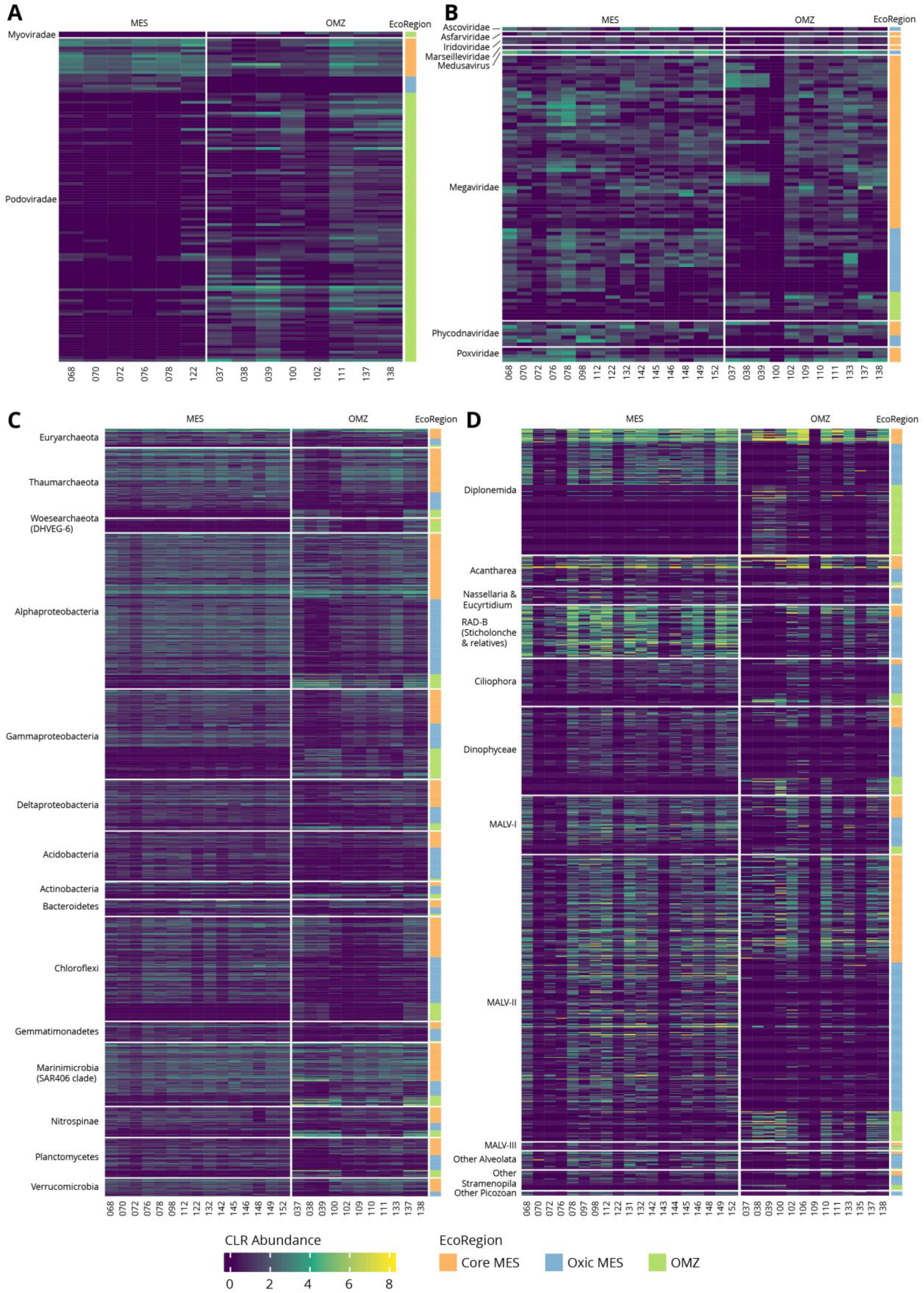
Heatmaps occurrence of OTUs assigned to mesopelagic eco-regions. A) Phages, B) NCLDV, C) Prokaryotes and D) pico-Eukaryotes.

The great majority of phage taxa (93.51%) occurred at similar abundance in all eco-regions (ubiquitous). Surprisingly, we did not identify any OTUs assigned specifically to an EPI eco-region, meaning that almost all taxa observed in the epipelagic layer were also present in the mesopelagic layer (Supplementary Figures 3, 4). This observation supports the seed-bank hypothesis raised by Brum et al. [60], and the correlation to the sinking particles observed here. On the other hand, we detected taxa specific to the mesopelagic layer, mostly related to the OMZ eco-region (Figure 4a, Supplementary Figures 3-4). This mesopelagic specificity agrees with the sharp increase in marine phage micro-diversity following depth, as previously shown by Gregory et al. [6]. Our results emphasize that one cause for phage stratification in the water column might be the adaptation to the mesopelagic environment. Two hypotheses arise here, 1) the environment acts as a strong driver, directly selecting phages independently of their hosts, and 2) there is higher phage-host specificity in the mesopelagic layer, promoting phage selection. Following the first hypothesis, we can posit that the environment can directly impact phage assemblage composition. The direct contact with the environment of free phage entities (released from their hosts) may reduce infectivity, degrade, or remove virus particles, and adversely affect adsorption to the host [69]. This direct environmental effect over marine phages was reported for different ionic gradients [70], daylight conditions, and temperature [71]. However, the enrichment of prokaryotic OTUs specific to mesopelagic regions (Figure 4, Supplementary Figures 3, 6), especially in OMZs, does not exclude the phage-host indirect selection relationship.

We found 136 mesopelagic-specific NCLDV OTUs out of 5538 in both oxic and OMZ eco-regions. Even though it represents a small number, these OTUs were highly abundant (Figure 4b, Supplementary Figures 3, 5). Most of the mesopelagic-specific NCLDV OTUs corresponded to the Core MES group (OMZ = 18 OTUs, Oxic MES = 31 OTUs, Core MES = 87 OTUs-Supplementary Material Table S3). NCLDV can encode genes such as transporters for ammonium, magnesium, and phosphate that are important in marine oligotrophic areas [72]. This characteristic can improve the host’s fitness in the short-term and ultimately favor NCLDV fecundity and endurance. This property is named NCLDV-mediated host reprogramming [72]. Our results therefore indicate that these entities are less diverse in mesopelagic waters and may successfully infect a wide range of hosts adapted to different oxygen concentrations.

Among the planktonic microorganisms, prokaryotes have been, so far, the most investigated group in OMZ regions, especially in the Pacific Ocean [20,21]. We could better distinguish the prokaryotic mesopelagic signatures between Oxic MES and OMZ, confirming the influence of oxygen reported here and in previous studies [20–22] (Figure 4c, Supplementary Figures 3, 6). We observed similar occurrences and abundances for the OMZ signature taxa in the Indian Ocean stations (IO - 037, 038, 039) and in stations PO - 100, 137, and 138 from the Pacific Ocean (Figure 4c). These Pacific stations are located in the open ocean (PO - 137 and 138 located in the Equatorial upwelling zone and station PO - 100 in the South Pacific Subtropical Gyre). They present a strong upwelling signature, disclosing an intense decrease in oxygen concentration almost reaching shallow waters. Likewise, the sampling stations in the Indian Ocean are located in well-stratified waters, markedly characterized by the abrupt decrease of oxygen concentration below the thermohaline at 100-120m depth, especially for stations 038 and 039. At the mesopelagic layer of the Indian Ocean stations, the oxygen concentration ranges from 0.83 to 3 μmol/kg, characterizing functionally anoxic waters since aerobic metabolisms cannot be sustained at this oxygen level [73]. The other OMZ stations in the Pacific Ocean (PO - 102, 109, 110, 111) are located in coastal areas. Although they are also under the influence of upwellings, with low oxygen content, the oxygen level does not correspond to anoxic conditions, so they are classified as suboxic waters. This microoxic condition of this environment is sufficient to completely alter the microbial metabolism delineating the community composition in these sites. In addition, differences in the formation of offshore and coastal upwelling, for instance, or the influence of river runoffs, transporting anthropogenic nutrient enrichment from the continent to coastal areas [73], could be crucial in supporting the differences we observed in OMZ communities.

The same clear enrichment in both OMZ anoxic and suboxic samples was observed for the pico-eukaryotic groups Diplonemida, MALV-II, and Dinophyceae, suggesting these OTUs as the true OMZ eukaryotic signatures (Figure 4d). Some OTUs of these groups exhibited similar occurrences in the anoxic Indian and Pacific Oceans but not in suboxic samples from the Pacific Ocean. However, we observed a lower number of pico-eukaryotic taxa in the OMZ eco-region, the prevailing OTUs being specific to Oxic MES locations in most cases.

Another step to better understand mesopelagic community dynamics is to dissect the ecological relationships among species that thrive in this layer. Co-occurrence networks can indicate how the environment may structure the community acting as a filter for resident species [74]. They can also give us glimpses of organisms’ potential ecological interactions based on species connectivity [74,75]. Combining the NCLDV, prokaryote, and pico-eukaryote data, we inferred a network containing 6,154 nodes and 12,935 edges (Figure 5a, Table 3). Due to the lower number of stations sampled for phages, we excluded this group from the analysis. We found mainly positive relationships (94%), suggesting a predominance of putative biotic interactions (e.g. competition, symbiosis) rather than taxa avoidance or exclusion. This dominance of positive relations was also reported for epipelagic plankton communities [36,76]. The global network had a modularity value greater than 0.4 (Table 3), indicating that the network has a modular structure [73]. Applying a community detection algorithm [57] on this global graph, we were able to delineate 36 distinct modules (or subnetworks), presumably corresponding to ecological communities. Three of them were mainly composed of OTUs significantly enriched in mesopelagic OTUs (Oxic MES enriched module 1, p-value.bh = 1.29e^-153^ and OMZ enriched modules 4, p-value.bh = 1.90e^-51^, and 17, p-value.bh = 2.1e^-13^; Figure 5). Together, these three modules covered almost the total richness found in the mesopelagic zone (Figure 5b), and presented similar values for the average degree, clustering coefficient, and average path length (Table 3). These parameters indicate a network complexity [74], hinting at distinct ecological niches within the mesopelagic layer.

**Figure 5:**
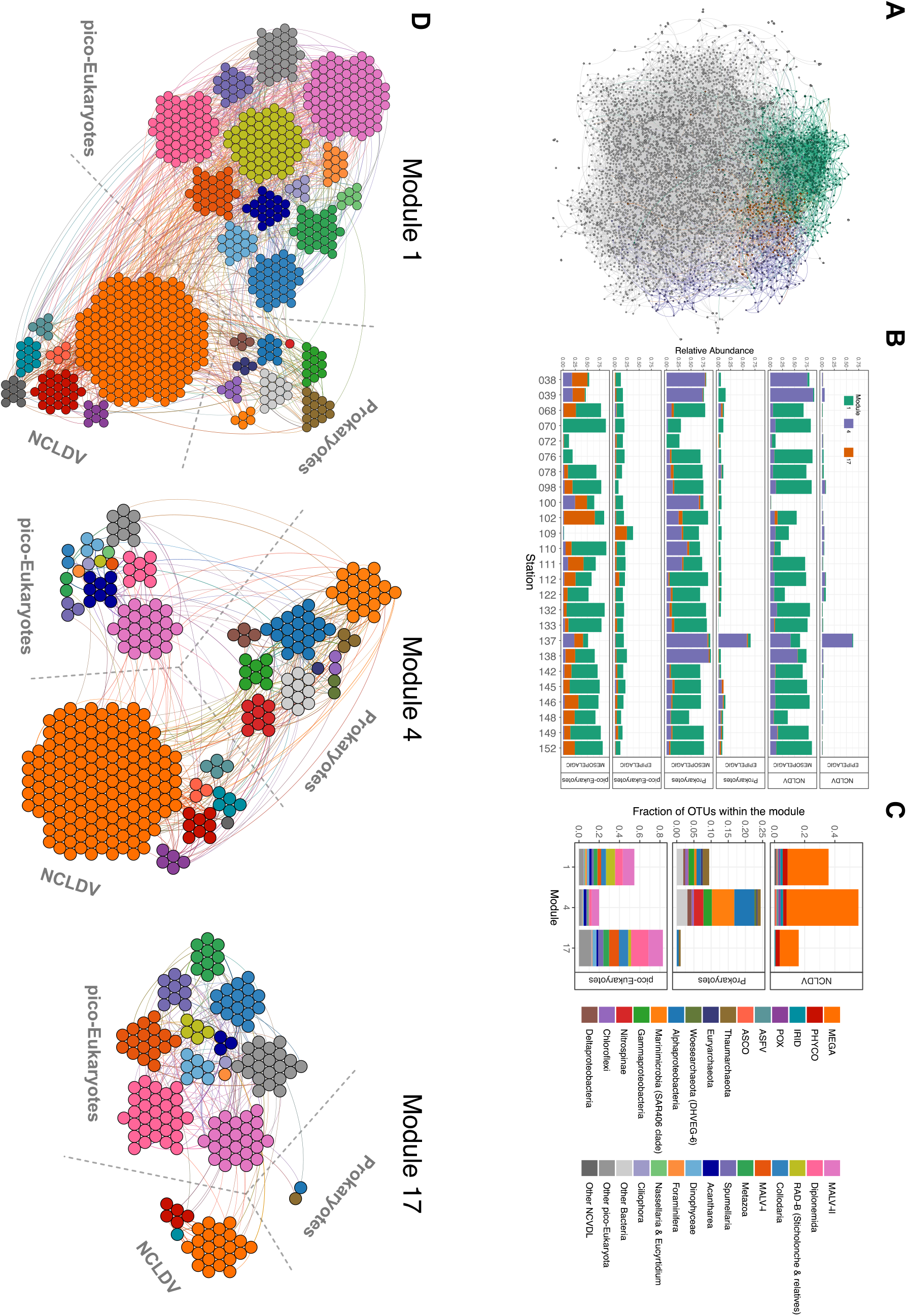
Co-occurrence network in epipelagic and mesopelagic communities. A) Global network, with connected modules for OMZ (purple and orange) and MES (green) highlighted. B) Relative taxa abundance in each module in each station and depth. C) Relative number of OTUs classified in taxonomic groups. D) Network representation of modules.

**Table 3.** Network topological features derived from global analysis including NCDLV, prokaryotes and pico-Eukaryotes samples in epipelagic and mesopelagic depths

| Parameter | Global | Mod 1 | Mod 4 | Mod 17 |
| --- | --- | --- | --- | --- |
| Nodes | 6154 | 731 | 323 | 175 |
| Positive edges | 12193 | 1236 | 480 | 223 |
| Negative edges | 742 | 70 | 49 | 9 |
| Av. Degree | 4.20 | 3.57 | 3.28 | 2.65 |
| Clustering | 0.03 | 0.03 | 0.09 | 0.05 |
| Density | 0.00 | 0.00 | 0.01 | 0.02 |
| Average path length | 7.28 | 6.01 | 6.27 | 6.30 |
| Betweenness | 0.01 | 0.05 | 0.10 | 0.22 |
| Degree Centralization | 0.00 | 0.01 | 0.02 | 0.04 |
| Modularity | 0.47 | 0.60 | 0.67 | 0.66 |

More precisely, the OMZ modules were composed of a few connected nodes (323 and 175 nodes for modules 4 and 17, respectively - Table 3), potentially indicating two distinct OMZ community niches. The Oxic MES module 1 counted more nodes composing the network associations (731 nodes), and all modules presented a variation in taxonomic composition and proportions. OMZ module 4 (OMZ-4) contained mainly prokaryotic (23%) and NCLDV (55%) OTUs (Figure 5c, d). Among the prokaryotes, we detected taxa previously determined as representing OMZ signatures (Nitrospinae, Marinimicrobia SAR 406, and Planctomycetes). As this module contains both prokaryotes and NCLDV OTUs, it suggests the existence of a confounding factor not captured in the dataset (such as large eukaryotes), as NCLDV are known to be specific to eukaryotes. Nevertheless, this correlation has been observed independently in another study [12]. On the other hand, module 17 (OMZ-17) is mainly composed of pico-eukaryotes OTUs (83%) (notably MALV-II (14%) and Diplonemida (17%) previously indicated as OMZ signatures) and NCLDV (16%) as expected due to virus-host relationships. Module 1 is taxonomically more diverse but consisted mainly of NCLDV and pico-eukaryotes. These groups accounted for 36% and 55%, respectively, of OTUs in this module (Figure 5c, d). NCLDV contributed to 598 associations (edges) in mesopelagic module 1, of which 177 occurred between NCLDV and pico-eukaryotes. NCLDV from the *Mimiviridae* family are the most numerous taxa in all three mesopelagic modules. *Mimiviridae* is a very abundant family in the ocean, present in various size ranges from piconanoplankton (0.8-5 μm) up to mesoplankton (180- 2,000 μm) [9,75]. This observation supports our finding that NCLDV are a prosperous group in mesopelagic waters, undertaking different strategies to endure in such environmental conditions. In all three modules, we observed the presence of Foraminifera, of which some species can use nitrate over oxygen as an electron acceptor, favoring their survival in OMZ regions [77].

Our converging results suggest that the mesopelagic zone can be characterized by at least three well-defined ecological niches (Oxic MES, OMZ-4 and OMZ-17), with established conditions and resources (abiotic and biotic) that allow the survival of a specific communities in these environments. Observed differences between OMZ and Oxic MES networks suggest a potential loss of connections and interactions among mesopelagic community members, directly affecting ecosystem stability due to habitat change.

### Conclusions

In this study, we explored mesopelagic pico-plankton ecological structuring and concluded that this component of oceanic plankton is heterogeneous with respect to environmental conditions. We could pinpoint the relevance of oxygen for all assemblages and the relation of particle flux with phages, NCLDV and pico-eukaryotes. These results reinforce the need to better understand the mesopelagic ecosystem in order to improve our comprehension of carbon export through the biological carbon pump in the twilight zone. Also, we show that intermediated water masses defined by their T/S profiles can explain the differences in the observed mesopelagic pico-plankton structure, pointing to the role of a set of environmental parameters for community composition.

By establishing eco-regions (Epipelagic, Oxic MES, and OMZ), we were able to discriminate specific mesopelagic signatures OTUs across all Life’s domains. While we recovered known markers for Oxic MES and OMZ regions at high taxonomic levels, we also found that most of these OTU signatures are observed at low taxonomic levels, which sometimes cannot be resolved using existing databases. Combining these OTU profiles within co-occurrence networks, we proposed three niches with biotic and abiotic conditions that appear to characterize mesopelagic ecosystems.

Limited access to data is usually the bottleneck for knowledge of mesopelagic dynamics. Our study benefits from a larger number of organism samples and distinct oceanic provinces. This allowed us to integrate these data and thus obtain an expanded vision of mesopelagic community structure and dynamics. Our results emphasize the need for a better understanding of mesopelagic life, particularly by improving our knowledge of oxic and oxygen-low mesopelagic-dwelling communities. This effort is especially necessary as climate change can be expected to expand marine OMZs in the future.

## Acknowledgments

This study is part of the “Ocean Plankton, Climate and Development” project conducted by the Tara Ocean Foundation with the support of the French Facility for the Global Environment (FFEM). Rigonato J., Budinich M., Murillo A.A., Brandão M.C., Pierella Karlusich J.J. and Soviadan Y.D. received financial support from FFEM to execute the project. Brandão, M.C. also received financial support from Coordination for the Improvement of Higher Education Personnel of Brazil (CAPES 99999.000487/2016-03). Budinich M. and Chaffron S. received financial support from the H2020 European Commission project AtlantECO (award number 862923). Tara Oceans (which includes both the Tara Oceans and Tara Oceans Polar Circle expeditions) would not exist without the leadership of the *Tara* Ocean Foundation and the continuous support of 23 institutes (https://fondationtaraocean.org). We further thank the commitment of the following sponsors: CNRS (in particular Groupement de Recherche GDR3280 and the Research Federation for the study of Global Ocean Systems Ecology and Evolution, FR2022/Tara Oceans-GOSEE), European Molecular Biology Laboratory (EMBL), Genoscope/CEA, The French Ministry of Research, and the French Government ‘Investissements d’Avenir’ programs OCEANOMICS (ANR-11-BTBR-0008), FRANCE GENOMIQUE (ANR-10-INBS-09-08), MEMO LIFE (ANR-10-LABX-54), CLIMACLOCK (ANR-20-CE20-0024) and PSL* Research University (ANR-11-IDEX-0001-02). This study benefits from the European Union’s Horizon 2020 Blue Growth research and innovation programme under grant agreement number 862923 (project AtlantECO). We also thank the support and commitment of agnès b. and Etienne Bourgois, the Prince Albert II de Monaco Foundation, the Veolia Foundation, Region Bretagne, Lorient Agglomeration, Serge Ferrari, World Courier. The global sampling effort was enabled by countless scientists and crew who sampled aboard the schooner *Tara* from 2009-2013, and we thank MERCATOR-CORIOLIS and ACRIST for providing daily satellite data during the expeditions. We are also grateful to the countries who graciously granted sampling permission. We want to thank to the reviewers and Editor for all the critics and suggestions that certainly improved the quality of the manuscript. We also acknowledge Noan Le Bescot (Ternog Design) for assistance in preparing figures. The authors declare that all data reported herein are fully and freely available from the date of publication, with no restrictions, and that all of the analyses, publications, and ownership of data are free from legal entanglement or restriction by the various nations whose waters the Tara Oceans expeditions sampled in. This article is contribution number 145 of Tara Oceans.

Tara Oceans Coordinators (alphabetical order) Silvia G. Acinas, Marcel Babin, Peer Bork, Emmanuel Boss, Chris Bowler, Guy Cochrane, Colomban de Vargas, Michael Follows, Gabriel Gorsky, Nigel Grimsley, Lionel Guidi, Pascal Hingamp, Daniele Iudicone, Olivier Jaillon, Stefanie Kandels, Lee Karp-Boss, Eric Karsenti, Fabrice Not, Hiroyuki Ogata, Stéphane Pesant, Nicole Poulton, Jeroen Raes, Christian Sardet, Sabrina Speich, Lars Stemmann, Matthew B. Sullivan, Shinichi Sunagawa and Patrick Wincker.

## Conflict of Interest

The authors declare that they have no conflict of interest.

---

**Supplementary Figure S1:**
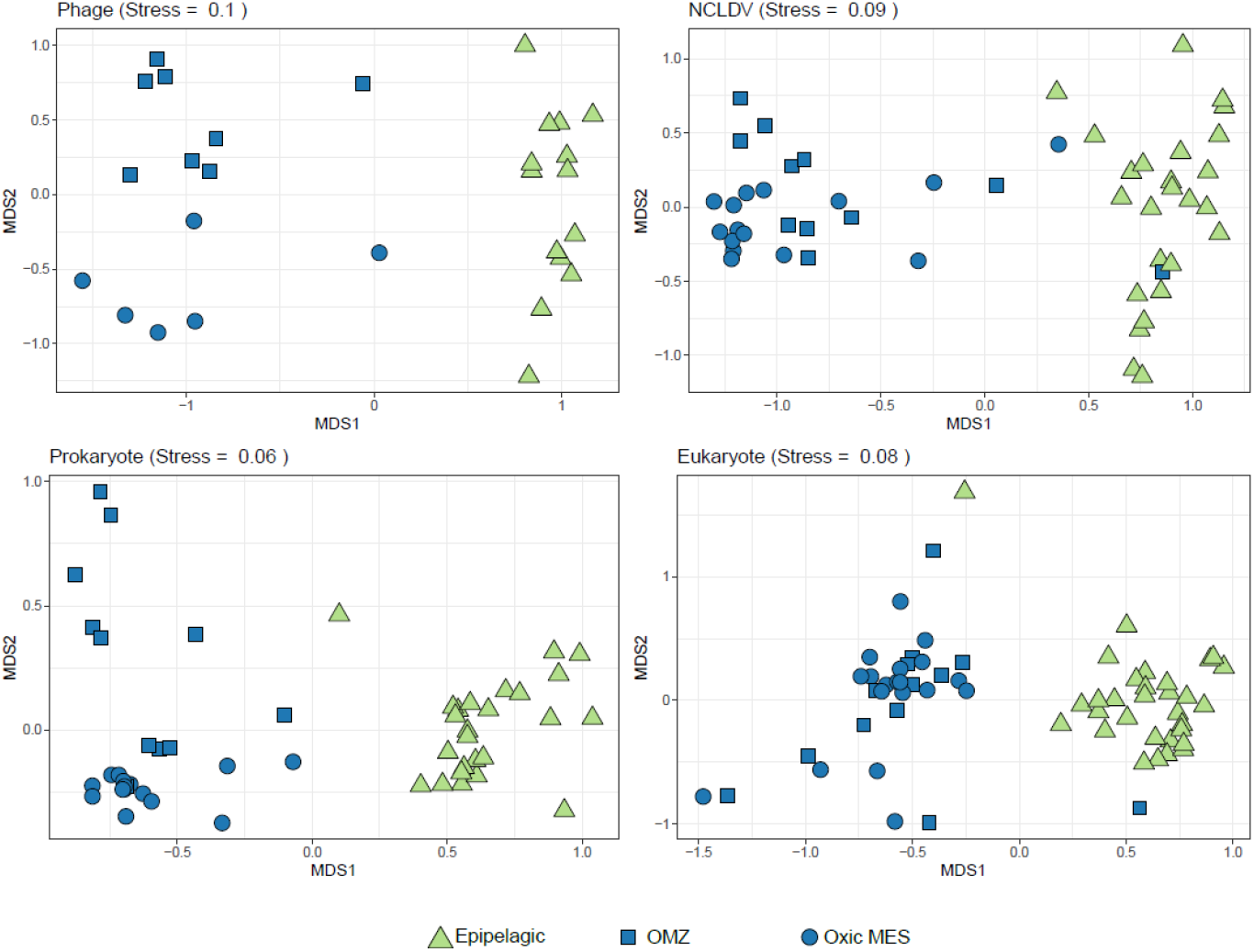
Non-metric multidimensional scaling (NMDS) showing epipelagic and mesopelagic community stratification for each organism group.

**Supplementary Figure S2:**
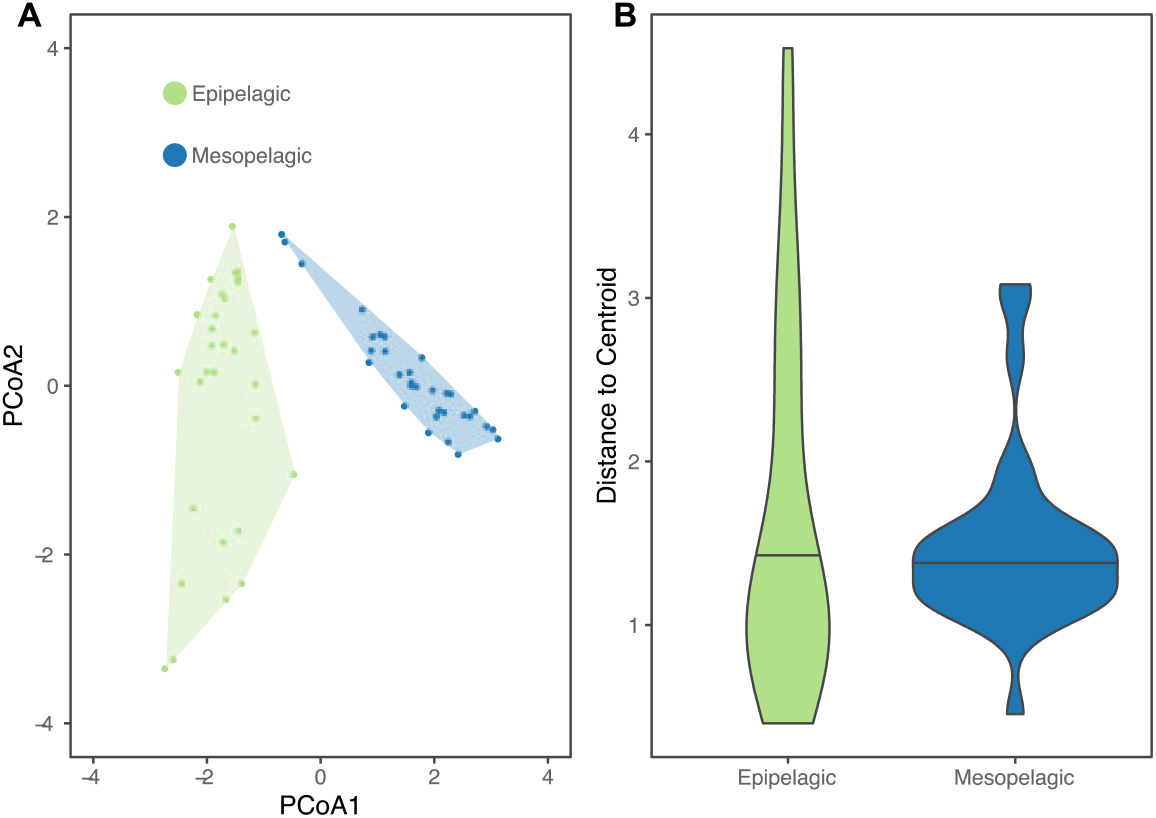
Epipelagic and mesopelagic group dispersion based on physical-chemical oceanic properties (Euclidian method). A) First two axes of PCoA. B) Dispersion of distances from samples to centroids.

**Supplementary Figure S3:**
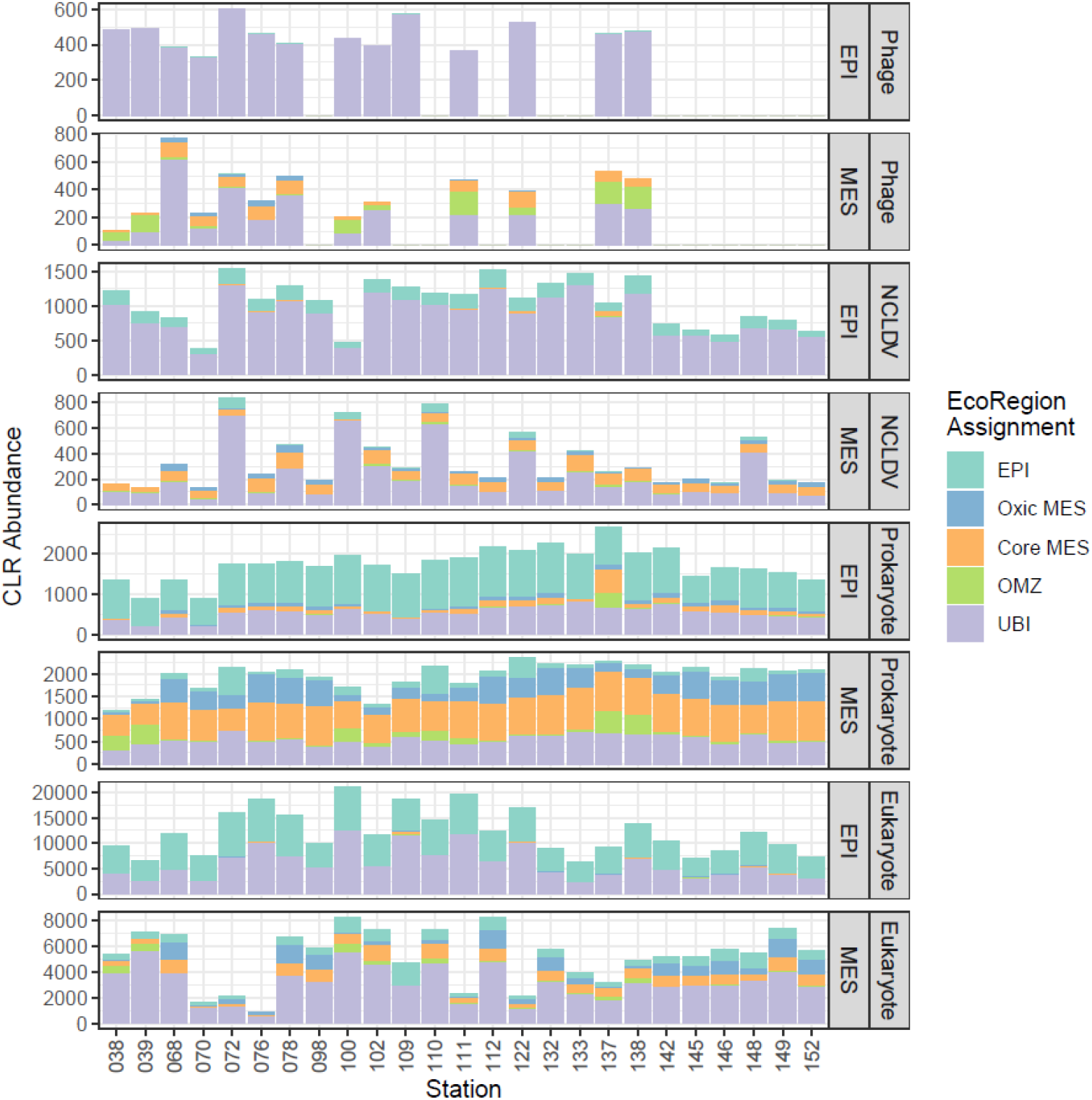
Occurrence of OTUs classified into different eco-regions by ocean layers.

**Supplementary Figure S4:**
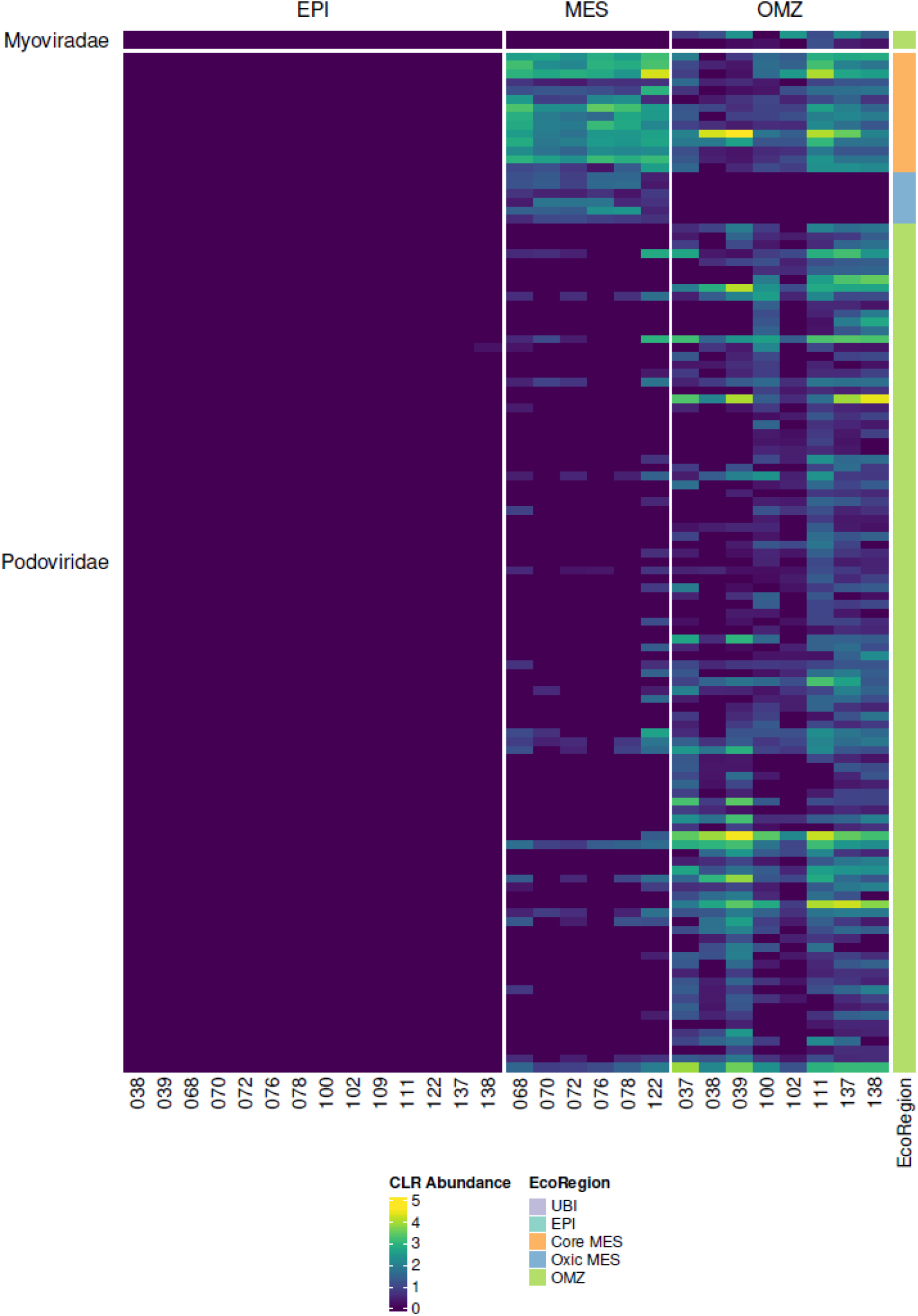
Normalized CLR abundance of phages and their preferred eco-region.

**Supplementary Figure S5:**
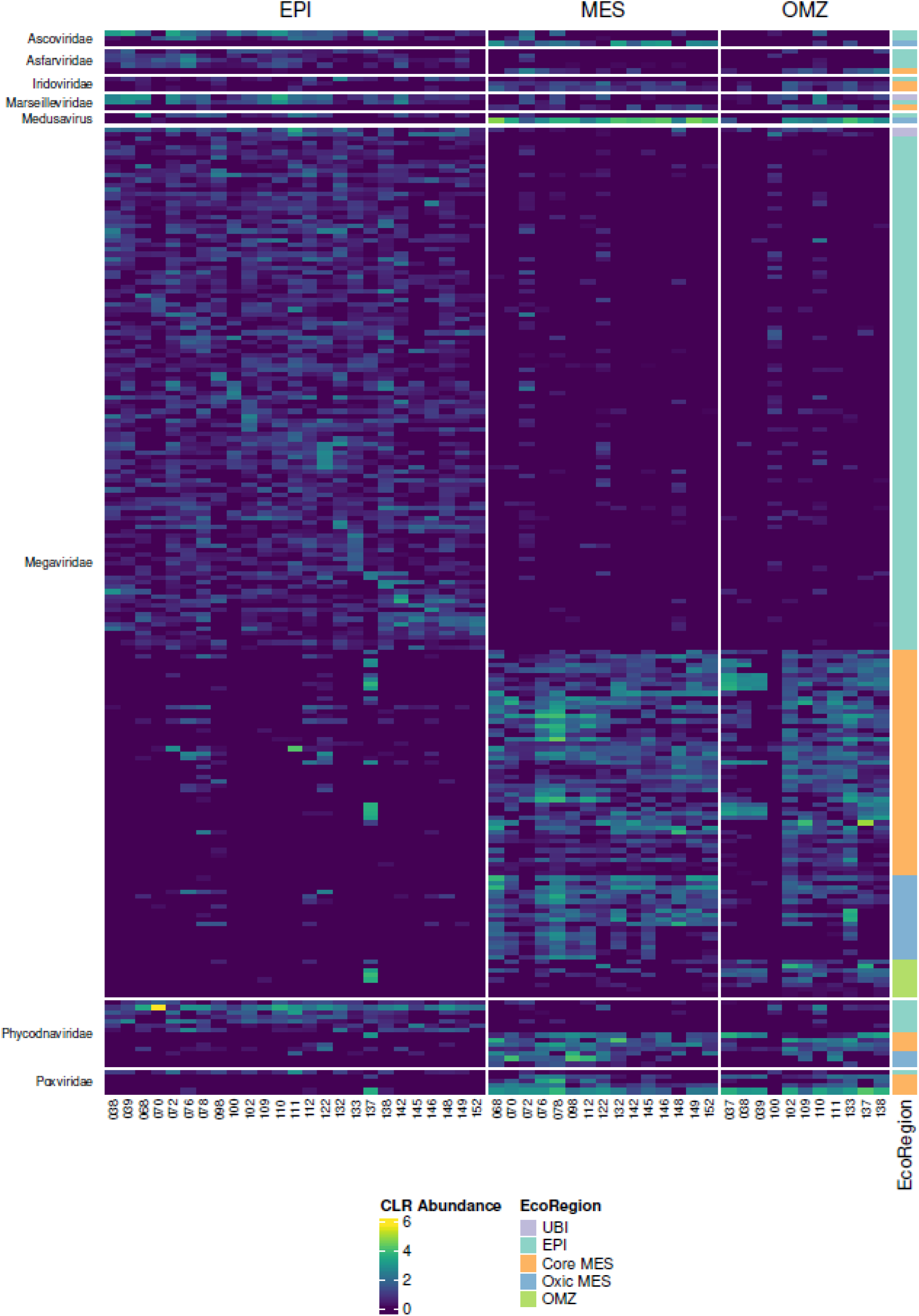
Normalized CLR abundance of NCLDV and their preferred eco-region.

**Supplementary Figure S6:**
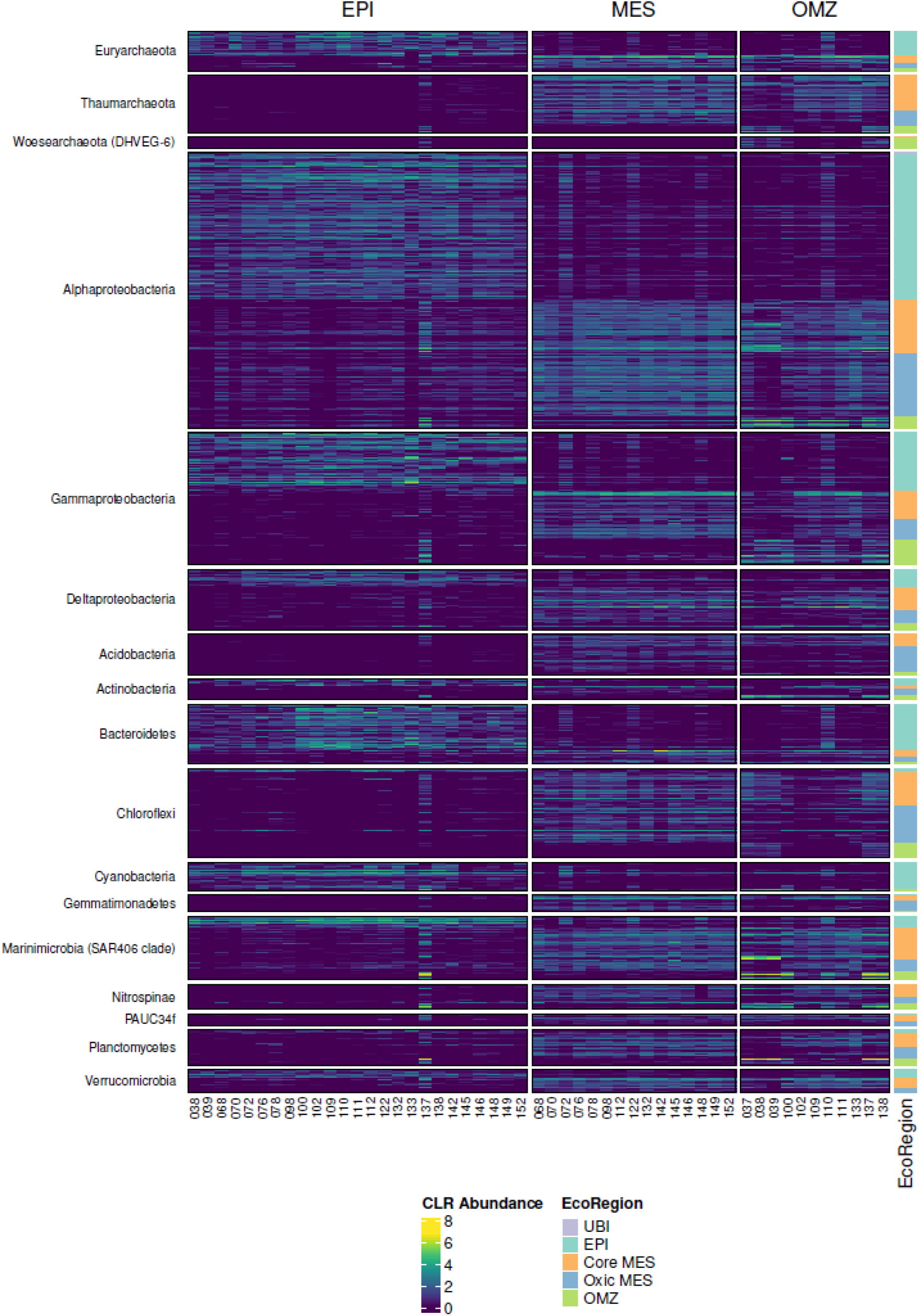
Normalized CLR abundance of prokaryotes and their preferred eco-region.

**Supplementary Figure S7:**
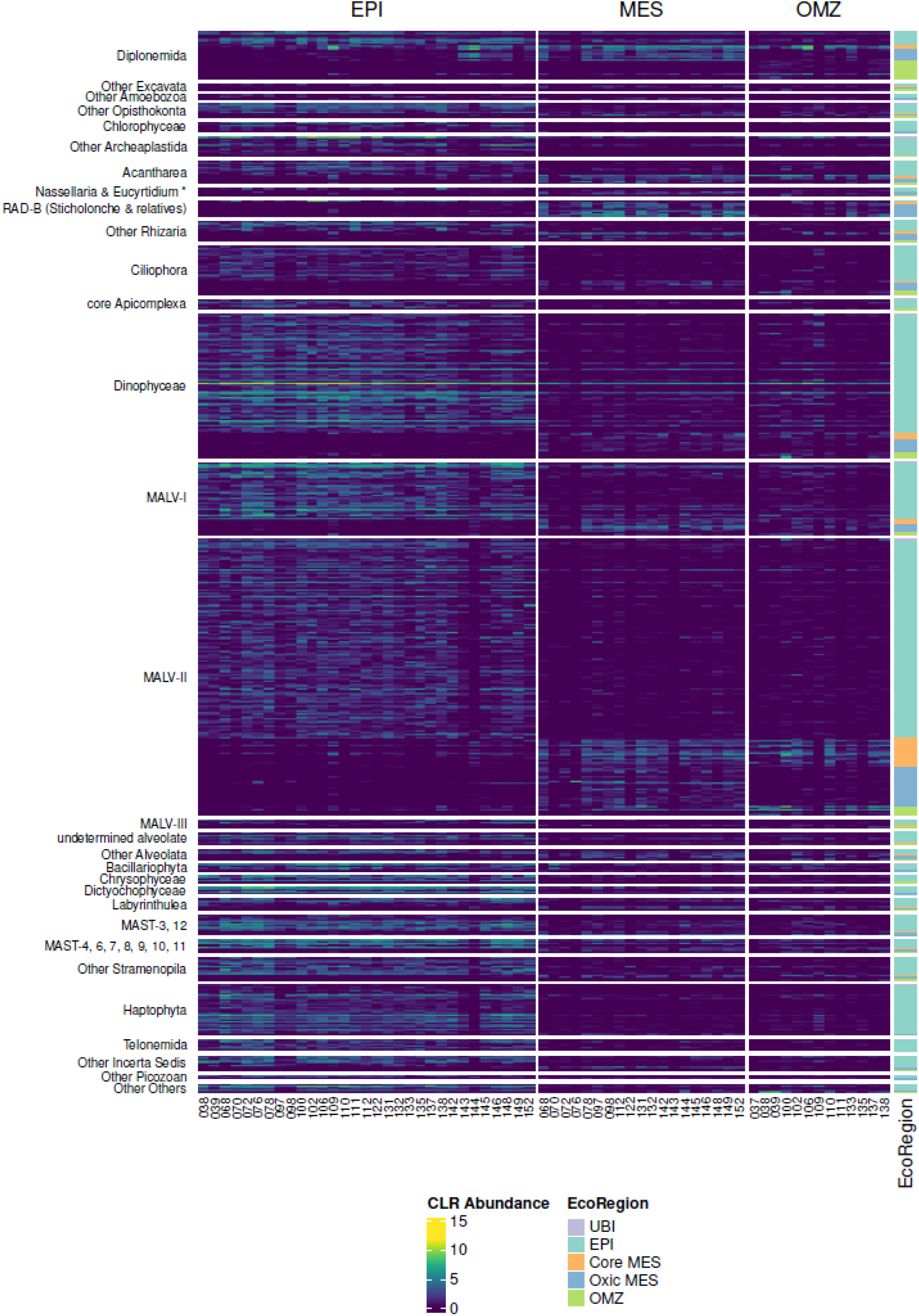
Normalized CLR abundance of pico-eukaryotes and their preferred eco-region.

## Supplementary Material and Methods

The present document details information about the *Tara* Oceans data availability and the pre-processing used for the analyses reported in Rigonato et al. 2021 “**Ocean-wide comparisons of mesopelagic planktonic community structures**”. Note that here we standardize the term “OTU” as “operational taxonomic unit” *sensu latu*, independent of clustering technique since taxonomy for the different assemblages was obtained using different methodologies. Also, it should be noted that keeping the original OTU definition datasets allowed direct comparison with previous work within the TARA Consortium.

## Data Processing

### Phage populations

Viral-phage contigs were assembled and identified from the 0.22 µm-filtered seawater viromes from the *Tara* Oceans expedition 2009-2013 and Malaspina 2010 global circumnavigation expedition using the methods described in (Gregory et al. *2019*). Prodigal 2.6.1 (Hyatt et al. 2010) was used to call ORFs (genes and proteins) using the ‘meta’ setting. The resulting proteins were clustered based on blast-defined sequence similarities and granularity of ‘2’ with MCL (Enright et al. 2002). The resulting genes were searched for four different primer sets for *gp23* (Table S2), the major capsid protein found in T4-related members of *Myoviridae*, and 10 different primer sets for *polA* (Table S2), the DNA polymerase found in *Podoviridae* (Adriaenssens et al. 2014). Because the genes are not always complete in metagenomic datasets, exact primer hits (100% nucleotide identity) of only the forward or the reverse primer were considered hits. Genes that hit these primers were translated into proteins and the MCL-defined protein clusters (PCs) that contained >=1 hit were extracted. PCs were annotated by running a combination of the reciprocal best blast hit analyses against the KEGG database (Kanehisa 2002) and blast against the UniProt Reference Clusters database (Suzek et al. 2007), searching for matches against the InterPro protein signature database using InterProScan (Zdobnov et al. 2001), and run hmmsearch against Pfam entries (Bateman et al. 2004) and then manually curated for *gp23* and *polA* genes, respectively. The *gp23* and *polA* proteins were matched to their gene sequences and dereplicated at 95% nucleotide identity across 100% the length into clusters using CD-HIT-EST (Li and Godzik 2006). The abundances of the *gp23* and *polA* clusters were calculated by pooling the abundances of the viral contigs containing the *gp23* or *polA* genes, respectively, per station.

In total, 848,507 viral contigs were identified. Viral contigs were grouped into populations if they shared ≥ 95% nucleotide identity across ≥ 80% of the genome (sensu) (Brum et al., 2015) using nucmer (Kurtz et al., 2004). With this, 488,130 total viral populations, here called OTUs, were obtained by the authors. For the present work, we used only *pol*B and *gp23* populations (9,048 OTUs).

**Table S2.**
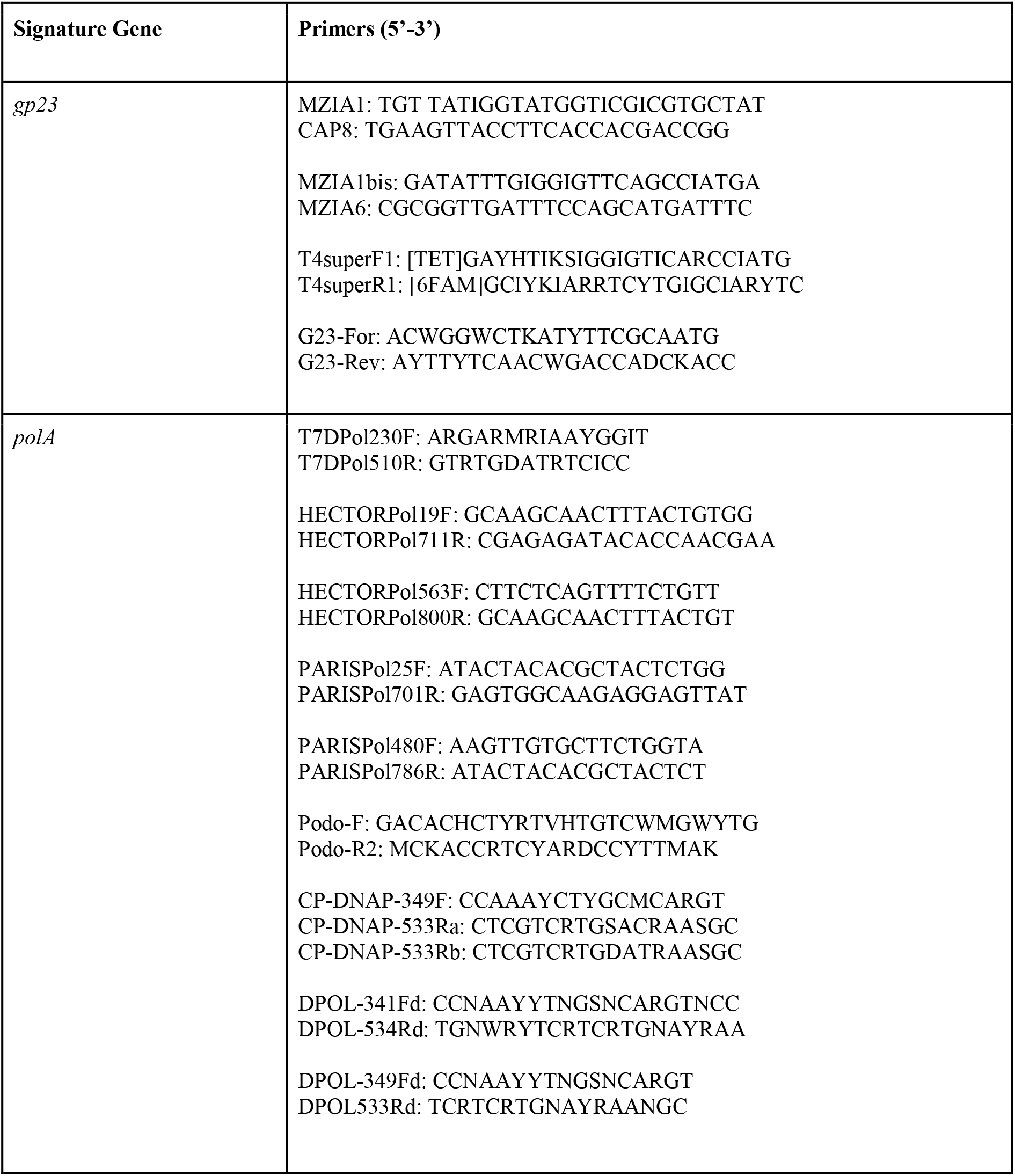
The *gp23* and *polA* primer sets used to search viral gene sequences

### Nucleo-Cytoplasmic Large DNA Virus - NCLDV

NCLDV sequence data was obtained from Endo et al., 2020. First, to assess the community composition of NCLDVs, *polB* was used as a marker gene for NCLDVs. 29,315 PolB sequences were obtained from the OM-RGC.v2 by using an in-house profile hidden Markov model of NCLDV *polB* sequences using the software HMMER, hmmsearch (version 3.1) with a threshold e-value of < 1 × 10^−5^.

To filter NCLDV derived sequences from those of other domains of life a phylogenetic tree was built based on 211 PolB reference protein sequences (eukaryotes, bacteria, archaea, phages, and NCLDVs). This tree included sequences from eight proposed families of NCLDVs: *Mimiviridae* (synonymous with *Megaviridae*), *Phycodnaviridae*, *Pithoviridae*, *Marseilleviridae*, *Ascoviridae*, *Iridoviridae*, *Asfarviridae*, and *Poxviridae*, and a sequence from *Medusavirus*, a novel NCLDV clade. The MAFFT-linsi (Kato and Standley 2013) was used for alignment and RAxML for maximum likelihood tree inference (Stamatakis 2006). NCLDV sequences derived from OM-RGC.v2 were aligned against the reference alignment using the MAFFT ‘addfragments’ option and then placed in the tree using the pplacer software (Matsen et al. 2010).

Next, NCLDV genes from the OM-RGC.v2 were used to obtain abundance profiles. Only samples from the pico- (0.22–1.6 μm or 0.22–3.0 μm) and femto- (<0.22 μm) size fractions were included. The sum of length-normalized *pol*B abundances ranged from 5.3 to 22,847.5 across samples, so the samples for which the sum of length-normalized *pol*B abundance was less than 50 (set as a proxy for low NCLDV frequency) were removed from the analysis. The abundance matrix was then standardized by the sample with the lowest sum of length-normalized *polB* abundance values.

### 16S rRNA gene mitags

The 16S rRNA gene dataset used in this manuscript was derived from environmental metagenomes following the approach of Logares et al. 2014, named “mitags”, and thoroughly described in Sunagawa et al. 2015 and Salazar et al. 2019. This approach is considered a powerful alternative to 16S rRNA gene amplicons as it overcomes PCR biases related to amplification and primer mismatch (Logares et al., 2014).

Briefly, 16S rRNA gene reference sequences from the SILVA database (Quast et al., 2013) were clustered at 97% sequence identity to balance the unequal taxa representation and define OTUs at the genus level. Then, the recruited mitags were mapped to cluster centroids of taxonomically annotated SILVA 16S reference sequences using USEARCH v9.2.64 (Edgar, 2010). Only the mitags mapping to a unique reference sequence were used to compute abundances at the OTU level. Mitags that mapped to more than one reference (OTU) were processed at a higher taxonomic level (domain, phylum, class, order, family or genus) common to all mapped OTUs. The abundance OTU table was generated by counting the number of mitags assigned to each taxon in each sample and the number of unassigned mitags. In total, 23,986 mitags were identified.

### 18S rRNA gene Metabarcoding

For 18S rRNA gene OTUs, conventional PCR marker amplification was used to investigate the eukaryotic diversity in Tara Ocean samples from de Vargas et al. 2015 and Ibarbalz et al. 2019. To obtain these OTUs, the authors applied a fast clustering method called *swarm* (Mahé et al., 2015), which can deliver high-resolution clusters, down to single-nucleotide differences with some additional post-processing, satisfying the ASV definition of Callahan et al. 2017. The *swarm* approach is free of arbitrary global clustering thresholds and input-order dependency that are the two main fundamental problems observed in traditional methods for clustering OTUs (Mahé et al., 2015).

The number of sequencing reads for each OTU was used as a proxy for abundance. OTUs were annotated using an in-house version of the PR2 database (Guillou et al. 2012, del Campo et al. 2018). In total, 474,303 OTUs were obtained.

A detailed explanation of read processing is available at http://taraoceans.sb-roscoff.fr/EukDiv/.

### Epipelagic Data Merging

Given the focus of the present study, we reclassified the samples as “Epipelagic” (SRF + DCM) and “Mesopelagic” (MES).

Here, we present results that support this decision, showing that SRF and DCM deviations are minor compared to MES samples. To combine both SRF and DCM samples, reads from the same OTUs at the same station were summed and then normalized by the total number of reads.

Figure 1 shows that the beta-diversity of the merged samples (EPI mod) do not differ from those obtained initially (*orig* SRF and DCM separately). The similarity between EPI and SRF or DCM was statistically confirmed by performing the ANOVA-like hypothesis test ANOSIM (Table S3; Clarke 1993). ANOSIM ranks the dissimilarity matrix to values between 0 and 1 (R values close to 0 suggest an even distribution and R values close to 1 suggest dissimilarity between groups) to compare the mean of ranked dissimilarities between groups to the mean of ranked dissimilarities within groups.

Figure 2 shows the dispersion of measures of the environmental samples recovered in situ during the biological sampling. We can observe that SFR and DCM are not statistically different.

**Figure 1.**
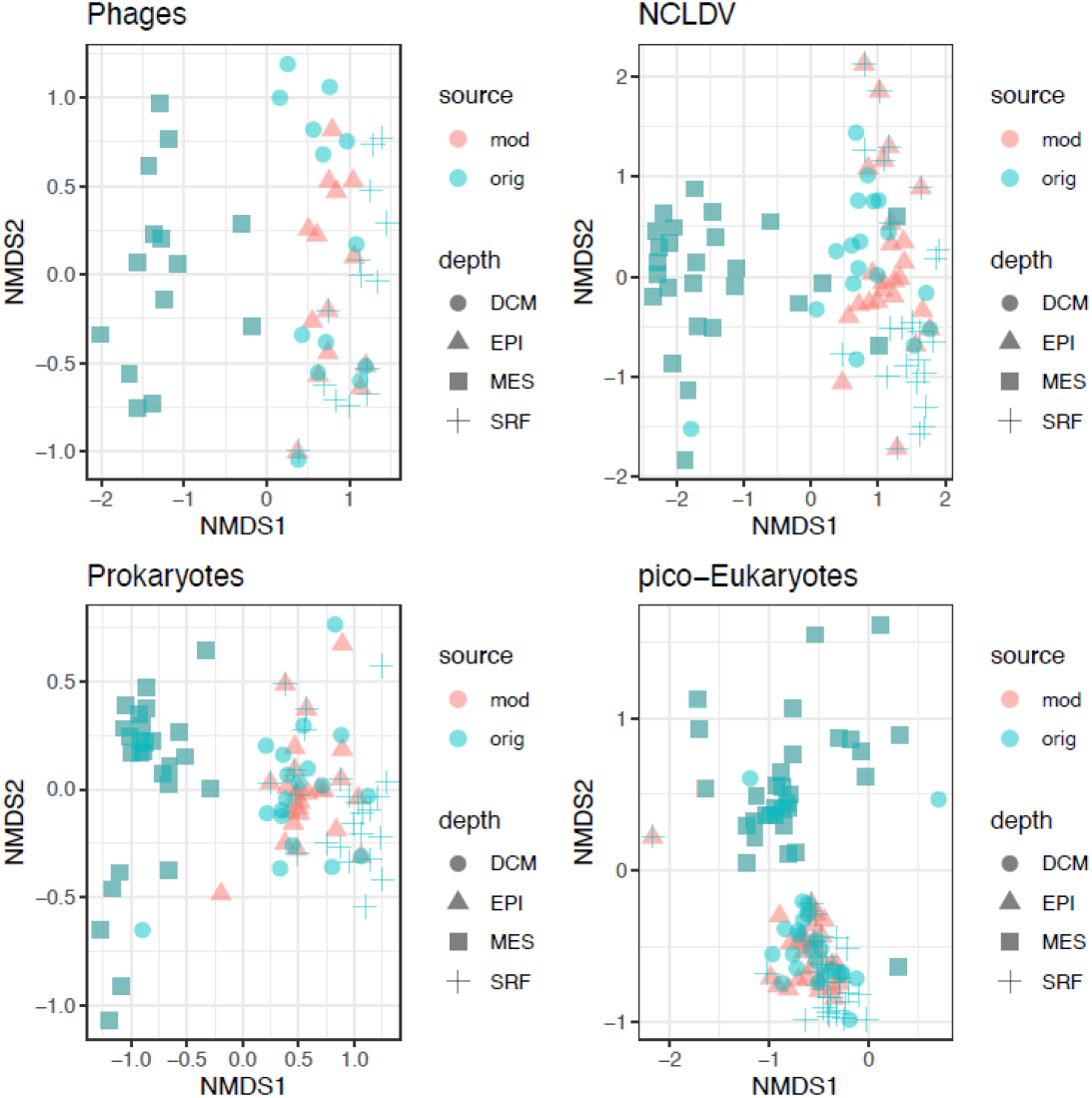
NMDS plots using the original datasets (*orig*, containing SRF, DCM, and MES layers) and the modified (*mod,* containing EPI and MES) datasets. MES samples are equal in both *mod* and *orig* datasets. Stress: 0.17 (virus), 0.14 (girus), 0.086 (Prokaryotes), 0.17 (Eukaryotes).

**Figure 2.**
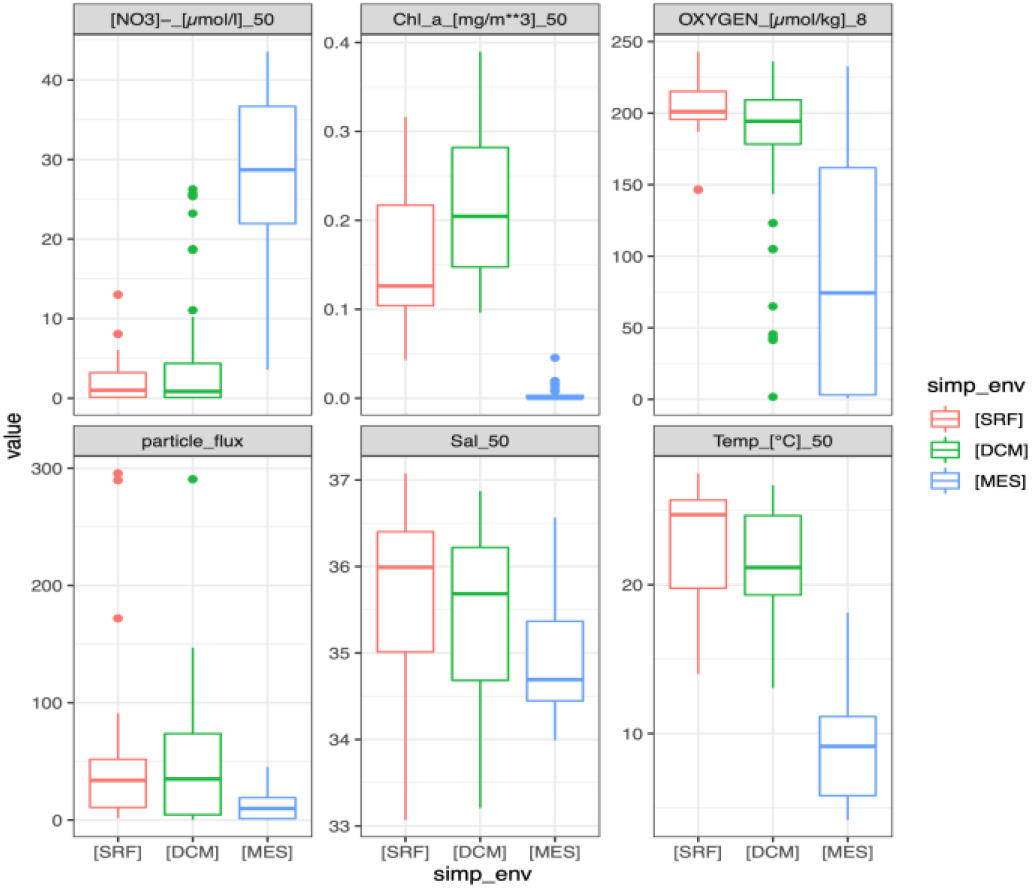
Environmental Parameters of Samples. An outlier in the particle flux (∼3428) was removed from the plot. Y-axis is in original parameter units from PANGEA (https://doi.pangaea.de/10.1594/PANGAEA.875582).

**Table S3.** ANOSIM statistics supporting the SRF and DCM data merging into EPI

| Assemblage | Depth1 | Depth2 | R | adj.p.val |
| --- | --- | --- | --- | --- |
| Phages | EPI | MES | 0,901 | 0,001 |
| Phages | EPI | DCM | 0,024 | 0,257 |
| Phages | EPI | SRF | 0,075 | 0,084 |
| Phages | MES | DCM | 0,883 | 0,001 |
| Phages | MES | SRF | 0,877 | 0,001 |
| Phages | DCM | SRF | 0,209 | 0,004 |
| NCLDV | EPI | MES | 0,772 | 0,001 |
| NCLDV | EPI | DCM | 0,045 | 0,118 |
| NCLDV | EPI | SRF | 0,028 | 0,136 |
| NCLDV | MES | DCM | 0,716 | 0,001 |
| NCLDV | MES | SRF | 0,815 | 0,001 |
| NCLDV | DCM | SRF | 0,096 | 0,020 |
| Prokaryotes | EPI | MES | 0,933 | 0,001 |
| Prokaryotes | EPI | DCM | 0,087 | 0,025 |
| Prokaryotes | EPI | SRF | 0,127 | 0,006 |
| Prokaryotes | MES | DCM | 0,855 | 0,001 |
| Prokaryotes | MES | SRF | 0,953 | 0,001 |
| Prokaryotes | DCM | SRF | 0,172 | 0,001 |
| pico-Eukaryotes | EPI | MES | 0,731 | 0,001 |
| pico-Eukaryotes | EPI | DCM | 0,140 | 0,002 |
| pico-Eukaryotes | EPI | SRF | 0,038 | 0,038 |
| pico-Eukaryotes | MES | DCM | 0,525 | 0,001 |
| pico-Eukaryotes | MES | SRF | 0,773 | 0,001 |
| pico-Eukaryotes | DCM | SRF | 0,179 | 0,001 |

## Notes

### Competing Interest Statement

The authors have declared no competing interest.

### Summary of Updates

The manuscript was revised, and new analysis was improved to clarify the results obtainable previously.

## References

1 Proud R, Cox MJ & Brierley AS (2017) Biogeography of the Global Ocean’s Mesopelagic Zone. Current Biology 27, 113–119.

2 Robinson C, Steinberg DK, Anderson TR, Arístegui J, Carlson CA, Frost JR, Ghiglione J-F, Hernández-León S, Jackson GA, Koppelmann R, Quéguiner B, Ragueneau O, Rassoulzadegan F, Robison BH, Tamburini C, Tanaka T, Wishner KF & Zhang J (2010) Mesopelagic zone ecology and biogeochemistry – a synthesis. Deep Sea Research Part II: Topical Studies in Oceanography 57, 1504–1518.

3 Mestre M, Ruiz-González C, Logares R, Duarte CM, Gasol JM & Sala MM (2018) Sinking particles promote vertical connectivity in the ocean microbiome. Proceedings of the National Academy of Sciences 115, E6799–E6807.

4 St. John MA, Borja A, Chust G, Heath M, Grigorov I, Mariani P, Martin AP & Santos RS (2016) A Dark Hole in Our Understanding of Marine Ecosystems and Their Services: Perspectives from the Mesopelagic Community. Frontiers in Marine Science 3.

5 Glover AG, Wiklund H, Chen C & Dahlgren TG (2018) Managing a sustainable deep-sea ‘blue economy’ requires knowledge of what actually lives there. eLife 7, e41319.

6 Gregory AC, Zayed AA, Conceição-Neto N, Temperton B, Bolduc B, Alberti A, Ardyna M, Arkhipova K, Carmichael M, Cruaud C, Dimier C, Domínguez-Huerta G, Ferland J, Kandels S, Liu Y, Marec C, Pesant S, Picheral M, Pisarev S, Poulain J, Tremblay JÉ, Vik D, Acinas SG, Babin M, Bork P, Boss E, Bowler C, Cochrane G, de Vargas C, Follows M, Gorsky G, Grimsley N, Guidi L, Hingamp P, Iudicone D, Jaillon O, Kandels-Lewis S, Karp-Boss L, Karsenti E, Not F, Ogata H, Poulton N, Raes J, Sardet C, Speich S, Stemmann L, Sullivan MB, Sunagawa S, Wincker P, Culley AI, Dutilh BE & Roux S (2019) Marine DNA Viral Macro- and Microdiversity from Pole to Pole. Cell 177, 1109–1123.e14.

7 Endo H, Blanc-Mathieu R, Li Y, Salazar G, Henry N, Labadie K, de Vargas C, Sullivan MB, Bowler C, Wincker P, Karp-Boss L, Sunagawa S & Ogata H (2020) Biogeography of marine giant viruses reveals their interplay with eukaryotes and ecological functions. Nature Ecology & Evolution.

8 Sunagawa S, Coelho LP, Chaffron S, Kultima JR, Labadie K, Salazar G, Djahanschiri B, Zeller G, Mende DR, Alberti A, Cornejo-Castillo FM, Costea PI, Cruaud C, D’Ovidio F, Engelen S, Ferrera I, Gasol JM, Guidi L, Hildebrand F, Kokoszka F, Lepoivre C, Lima-Mendez G, Poulain J, Poulos BT, Royo-Llonch M, Sarmento H, Vieira-Silva S, Dimier C, Picheral M, Searson S, Kandels-Lewis S, Bowler C, de Vargas C, Gorsky G, Grimsley N, Hingamp P, Iudicone D, Jaillon O, Not F, Ogata H, Pesant S, Speich S, Stemmann L, Sullivan MB, Weissenbach J, Wincker P, Karsenti E, Raes J, Acinas SG & Bork P (2015) Ocean plankton. Structure and function of the global ocean microbiome. *Science (New York*, NY) 348, 1261359.

9 Salazar G, Paoli L, Alberti A, Huerta-Cepas J, Ruscheweyh H-J, Cuenca M, Field CM, Coelho LP, Cruaud C, Engelen S, Gregory AC, Labadie K, Marec C, Pelletier E, Royo-Llonch M, Roux S, Sánchez P, Uehara H, Zayed AA, Zeller G, Carmichael M, Dimier C, Ferland J, Kandels S, Picheral M, Pisarev S, Poulain J, Acinas SG, Babin M, Bork P, Bowler C, de Vargas C, Guidi L, Hingamp P, Iudicone D, Karp-Boss L, Karsenti E, Ogata H, Pesant S, Speich S, Sullivan MB, Wincker P, Sunagawa S, Acinas SG, Babin M, Bork P, Boss E, Bowler C, Cochrane G, de Vargas C, Follows M, Gorsky G, Grimsley N, Guidi L, Hingamp P, Iudicone D, Jaillon O, Kandels-Lewis S, Karp-Boss L, Karsenti E, Not F, Ogata H, Pesant S, Poulton N, Raes J, Sardet C, Speich S, Stemmann L, Sullivan MB, Sunagawa S & Wincker P (2019) Gene Expression Changes and Community Turnover Differentially Shape the Global Ocean Metatranscriptome. Cell 179, 1068–1083.e21.

10 Giovannoni SJ & Vergin KL (2012) Seasonality in Ocean Microbial Communities. Science 335, 671–676.

11 Giner CR, Pernice MC, Balagué V, Duarte CM, Gasol JM, Logares R & Massana R (2020) Marked changes in diversity and relative activity of picoeukaryotes with depth in the world ocean. The ISME Journal 14, 437–449.

12 Ibarbalz FM, Henry N, Brandão MC, Martini S, Busseni G, Byrne H, Coelho LP, Endo H, Gasol JM, Gregory AC, Mahé F, Rigonato J, Royo-Llonch M, Salazar G, Sanz-Sáez I, Scalco E, Soviadan D, Zayed AA, Zingone A, Labadie K, Ferland J, Marec C, Kandels S, Picheral M, Dimier C, Poulain J, Pisarev S, Carmichael M, Pesant S, Babin M, Boss E, Iudicone D, Jaillon O, Acinas SG, Ogata H, Pelletier E, Stemmann L, Sullivan MB, Sunagawa S, Bopp L, de Vargas C, Karp-Boss L, Wincker P, Lombard F, Bowler C, Zinger L, Acinas SG, Babin M, Bork P, Boss E, Bowler C, Cochrane G, de Vargas C, Follows M, Gorsky G, Grimsley N, Guidi L, Hingamp P, Iudicone D, Jaillon O, Kandels S, Karp-Boss L, Karsenti E, Not F, Ogata H, Pesant S, Poulton N, Raes J, Sardet C, Speich S, Stemmann L, Sullivan MB, Sunagawa S & Wincker P (2019) Global Trends in Marine Plankton Diversity across Kingdoms of Life. Cell 179, 1084–1097.e21.

13 Baltar F, Arístegui J, Gasol JM & Herndl GJ (2012) Microbial Functioning and Community Structure Variability in the Mesopelagic and Epipelagic Waters of the Subtropical Northeast Atlantic Ocean. Appl Environ Microbiol 78, 3309–3316.

14 Guidi L, Chaffron S, Bittner L, Eveillard D, Larhlimi A, Roux S, Darzi Y, Audic S, Berline L, Brum JR, Coelho LP, Espinoza JCI, Malviya S, Sunagawa S, Dimier C, Kandels-Lewis S, Picheral M, Poulain J, Searson S, Stemmann L, Not F, Hingamp P, Speich S, Follows M, Karp-Boss L, Boss E, Ogata H, Pesant S, Weissenbach J, Wincker P, Acinas SG, Bork P, De Vargas C, Iudicone D, Sullivan MB, Raes J, Karsenti E, Bowler C & Gorsky G (2016) Plankton networks driving carbon export in the oligotrophic ocean. Nature 532, 465–470.

15 Landry Z, Swan BK, Herndl GJ, Stepanauskas R & Giovannoni SJ (2017) SAR202 Genomes from the Dark Ocean Predict Pathways for the Oxidation of Recalcitrant Dissolved Organic Matter. mBio 8.

16 Capone DG & Hutchins DA (2013) Microbial biogeochemistry of coastal upwelling regimes in a changing ocean. Nature Geoscience 6, 711–717.

17 Breitburg D, Levin LA, Oschlies A, Grégoire M, Chavez FP, Conley DJ, Garçon V, Gilbert D, Gutiérrez D, Isensee K, Jacinto GS, Limburg KE, Montes I, Naqvi SWA, Pitcher GC, Rabalais NN, Roman MR, Rose KA, Seibel BA, Telszewski M, Yasuhara M & Zhang J (2018) Declining oxygen in the global ocean and coastal waters. Science 359, eaam7240.

18 Stevens H & Ulloa O (2008) Bacterial diversity in the oxygen minimum zone of the eastern tropical South Pacific. Environmental Microbiology 10, 1244–1259.

19 Stewart FJ, Ulloa O & DeLong EF (2012) Microbial metatranscriptomics in a permanent marine oxygen minimum zone: OMZ community gene expression. Environmental Microbiology 14, 23–40.

20 Ulloa O, Wright JJ, Belmar L & Hallam SJ (2013) Pelagic Oxygen Minimum Zone Microbial Communities. In The Prokaryotes (Rosenberg E, DeLong EF, Lory S, Stackebrandt E, & Thompson F, eds), pp. 113–122. Springer Berlin Heidelberg, Berlin, Heidelberg.

21 Ulloa O & Pantoja S (2009) The oxygen minimum zone of the eastern South Pacific. Deep Sea Research Part II: Topical Studies in Oceanography 56, 987–991.

22 Wright JJ, Konwar KM & Hallam SJ (2012) Microbial ecology of expanding oxygen minimum zones. Nature Reviews Microbiology 10, 381–394.

23. Divya B (2017) Bacterial Community Profiling of the Arabian Sea Oxygen Minimum Zone Sediments using Cultivation Independent Approach. EIMBO 1.

24 Duret MT, Pachiadaki MG, Stewart FJ, Sarode N, Christaki U, Monchy S, Srivastava A & Edgcomb VP (2015) Size-fractionated diversity of eukaryotic microbial communities in the Eastern Tropical North Pacific oxygen minimum zone. FEMS Microbiology Ecology 91.

25 Orsi W, Song YC, Hallam S & Edgcomb V (2012) Effect of oxygen minimum zone formation on communities of marine protists. The ISME Journal 6, 1586–1601.

26 Parris DJ, Ganesh S, Edgcomb VP, DeLong EF & Stewart FJ (2014) Microbial eukaryote diversity in the marine oxygen minimum zone off northern Chile. Frontiers in Microbiology 5.

27 Vik D, Gazitúa MC, Sun CL, Zayed AA, Aldunate M, Mulholland MR, Ulloa O & Sullivan MB (2021) Genome-resolved viral ecology in a marine oxygen minimum zone. Environmental Microbiology 23, 2858–2874.

28 Gazitúa MC, Vik DR, Roux S, Gregory AC, Bolduc B, Widner B, Mulholland MR, Hallam SJ, Ulloa O & Sullivan MB (2021) Potential virus-mediated nitrogen cycling in oxygen-depleted oceanic waters. ISME J 15, 981–998.

29 Chow C-ET, Winget DM, White RA, Hallam SJ & Suttle CA (2015) Combining genomic sequencing methods to explore viral diversity and reveal potential virus-host interactions. Front Microbiol 6.

30. Rusch DB, Halpern AL, Sutton G, Heidelberg KB, Williamson S, Yooseph S, Wu D, Eisen JA, Hoffman JM, Remington K, Beeson K, Tran B, Smith H, Baden-Tillson H, Stewart C, Thorpe J, Freeman J, Andrews-Pfannkoch C, Venter JE, Li K, Kravitz S, Heidelberg JF, Utterback T, Rogers Y-H, Falcón LI, Souza V, Bonilla-Rosso G, Eguiarte LE, Karl DM, Sathyendranath S, Platt T, Bermingham E, Gallardo V, Tamayo-Castillo G, Ferrari MR, Strausberg RL, Nealson K, Friedman R, Frazier M & Venter JC (2007) The Sorcerer II Global Ocean Sampling Expedition: Northwest Atlantic through Eastern Tropical Pacific. PLOS Biology 5, e77.

31 Karsenti E, Acinas SG, Bork P, Bowler C, De Vargas C, Raes J, Sullivan M, Arendt D, Benzoni F, Claverie J-M, Follows M, Gorsky G, Hingamp P, Iudicone D, Jaillon O, Kandels-Lewis S, Krzic U, Not F, Ogata H, Pesant S, Reynaud EG, Sardet C, Sieracki ME, Speich S, Velayoudon D, Weissenbach J, Wincker P, & the Tara Oceans Consortium (2011) A Holistic Approach to Marine Eco-Systems Biology. PLoS Biology 9, e1001177.

32 Pernice MC, Forn I, Gomes A, Lara E, Alonso-Sáez L, Arrieta JM, del Carmen Garcia F, Hernando-Morales V, MacKenzie R, Mestre M, Sintes E, Teira E, Valencia J, Varela MM, Vaqué D, Duarte CM, Gasol JM & Massana R (2015) Global abundance of planktonic heterotrophic protists in the deep ocean. The ISME Journal 9, 782–792.

33 Morris RM, Rappé MS, Urbach E, Connon SA & Giovannoni SJ (2004) Prevalence of the *Chloroflexi* -Related SAR202 Bacterioplankton Cluster throughout the Mesopelagic Zone and Deep Ocean. Appl Environ Microbiol 70, 2836–2842.

34 Mehrshad M, Rodriguez-Valera F, Amoozegar MA, López-García P & Ghai R (2018) The enigmatic SAR202 cluster up close: shedding light on a globally distributed dark ocean lineage involved in sulfur cycling. The ISME Journal 12, 655–668.

35 Hidalgo M & Browman HI (2019) Developing the knowledge base needed to sustainably manage mesopelagic resources. ICES Journal of Marine Science 76, 609–615.

36 Lima-Mendez G, Faust K, Henry N, Decelle J, Colin S, Carcillo F, Chaffron S, Ignacio-Espinosa JC, Roux S, Vincent F, Bittner L, Darzi Y, Wang J, Audic S, Berline L, Bontempi G, Cabello AM, Coppola L, Cornejo-Castillo FM, d’Ovidio F, De Meester L, Ferrera I, Garet-Delmas M-J, Guidi L, Lara E, Pesant S, Royo-Llonch M, Salazar G, Sanchez P, Sebastian M, Souffreau C, Dimier C, Picheral M, Searson S, Kandels-Lewis S, Tara Oceans coordinators, Gorsky G, Not F, Ogata H, Speich S, Stemmann L, Weissenbach J, Wincker P, Acinas SG, Sunagawa S, Bork P, Sullivan MB, Karsenti E, Bowler C, de Vargas C & Raes J (2015) Determinants of community structure in the global plankton interactome. Science 348, 1262073–1262073.

37 Louca S, Hawley AK, Katsev S, Torres-Beltran M, Bhatia MP, Kheirandish S, Michiels CC, Capelle D, Lavik G, Doebeli M, Crowe SA & Hallam SJ (2016) Integrating biogeochemistry with multiomic sequence information in a model oxygen minimum zone. Proceedings of the National Academy of Sciences 113, E5925–E5933.

38 Sunagawa S, Acinas SG, Bork P, Bowler C, Acinas SG, Babin M, Bork P, Boss E, Bowler C, Cochrane G, de Vargas C, Follows M, Gorsky G, Grimsley N, Guidi L, Hingamp P, Iudicone D, Jaillon O, Kandels S, Karp-Boss L, Karsenti E, Lescot M, Not F, Ogata H, Pesant S, Poulton N, Raes J, Sardet C, Sieracki M, Speich S, Stemmann L, Sullivan MB, Sunagawa S, Wincker P, Eveillard D, Gorsky G, Guidi L, Iudicone D, Karsenti E, Lombard F, Ogata H, Pesant S, Sullivan MB, Wincker P & de Vargas C (2020) Tara Oceans: towards global ocean ecosystems biology. Nature Reviews Microbiology 18, 428–445.

39. Guidi L, Picheral M, Pesant S, Tara Oceans Consortium C & Tara Oceans Expedition P (2017) Environmental context of all samples from the Tara Oceans Expedition (2009-2013), about sensor data in the targeted environmental feature PANGAEA - Data Publisher for Earth & Environmental Science.

40. Guidi L, Ras J, Claustre H, Pesant S, Tara Oceans Consortium C & Tara Oceans Expedition P (2017) Environmental context of all samples from the Tara Oceans Expedition (2009-2013), about pigment concentrations (HPLC) in the targeted environmental feature PANGAEA - Data Publisher for Earth & Environmental Science.

41. Guidi L, Morin P, Coppola L, Tremblay J-É, Pesant S, Tara Oceans Consortium C & Tara Oceans Expedition P (2017) Environmental context of all samples from the Tara Oceans Expedition (2009-2013), about nutrients in the targeted environmental feature PANGAEA - Data Publisher for Earth & Environmental Science.

42. Pesant S, Tara Oceans Consortium C & Tara Oceans Expedition P (2017) Methodology used on board to prepare samples from the Tara Oceans Expedition (2009-2013) PANGAEA - Data Publisher for Earth & Environmental Science.

43. Speich S, Chaffron S, Ardyna M, Pesant S, Tara Oceans Consortium C & Tara Oceans Expedition P (2017) Environmental context of all samples from the Tara Oceans Expedition (2009-2013), about the water column features at the sampling location PANGAEA - Data Publisher for Earth & Environmental Science.

44 Picheral M, Guidi L, Stemmann L, Karl DM, Iddaoud G & Gorsky G (2010) The Underwater Vision Profiler 5: An advanced instrument for high spatial resolution studies of particle size spectra and zooplankton. Limnology and Oceanography: Methods 8, 462–473.

45. Picheral M, Searson S, Taillandier V, Bricaud A, Boss E, Ras J, Claustre H, Ouhssain M, Morin P, Tremblay J-É, Coppola L, Gattuso J-P, Metzl N, Thuillier D, Gorsky G, Tara Oceans Consortium C & Tara Oceans Expedition P (2014) Vertical profiles of environmental parameters measured on discrete water samples collected with Niskin bottles during the Tara Oceans expedition 2009-2013 PANGAEA - Data Publisher for Earth & Environmental Science.

46. 46 Picheral M, Colin S & Irisson JO (2017) EcoTaxa, A Tool for the Taxonomic Classification of Images.

47 Schlitzer R (2002) Interactive analysis and visualization of geoscience data with Ocean Data View. Computers & Geosciences 28, 1211–1218.

48 Pesant S, Not F, Picheral M, Kandels-Lewis S, Le Bescot N, Gorsky G, Iudicone D, Karsenti E, Speich S, Troublé R, Dimier C, Searson S, Acinas SG, Bork P, Boss E, Bowler C, De Vargas C, Follows M, Gorsky G, Grimsley N, Hingamp P, Iudicone D, Jaillon O, Kandels-Lewis S, Karp-Boss L, Karsenti E, Krzic U, Not F, Ogata H, Pesant S, Raes J, Reynaud EG, Sardet C, Sieracki M, Speich S, Stemmann L, Sullivan MB, Sunagawa S, Velayoudon D, Weissenbach J & Wincker P (2015) Open science resources for the discovery and analysis of Tara Oceans data. Scientific Data 2, 150023.

49 Alberti A, Poulain J, Engelen S, Labadie K, Romac S, Ferrera I, Albini G, Aury J-M, Belser C, Bertrand A, Cruaud C, Da Silva C, Dossat C, Gavory F, Gas S, Guy J, Haquelle M, Jacoby E, Jaillon O, Lemainque A, Pelletier E, Samson G, Wessner M, Bazire P, Beluche O, Bertrand L, Besnard-Gonnet M, Bordelais I, Boutard M, Dubois M, Dumont C, Ettedgui E, Fernandez P, Garcia E, Aiach NG, Guerin T, Hamon C, Brun E, Lebled S, Lenoble P, Louesse C, Mahieu E, Mairey B, Martins N, Megret C, Milani C, Muanga J, Orvain C, Payen E, Perroud P, Petit E, Robert D, Ronsin M, Vacherie B, Acinas SG, Royo-Llonch M, Cornejo-Castillo FM, Logares R, Fernández-Gómez B, Bowler C, Cochrane G, Amid C, Hoopen PT, De Vargas C, Grimsley N, Desgranges E, Kandels-Lewis S, Ogata H, Poulton N, Sieracki ME, Stepanauskas R, Sullivan MB, Brum JR, Duhaime MB, Poulos BT, Hurwitz BL, Acinas SG, Bork P, Boss E, Bowler C, De Vargas C, Follows M, Gorsky G, Grimsley N, Hingamp P, Iudicone D, Jaillon O, Kandels-Lewis S, Karp-Boss L, Karsenti E, Not F, Ogata H, Pesant S, Raes J, Sardet C, Sieracki ME, Speich S, Stemmann L, Sullivan MB, Sunagawa S, Wincker P, Pesant S, Karsenti E & Wincker P (2017) Viral to metazoan marine plankton nucleotide sequences from the Tara Oceans expedition. Scientific Data 4, 170093.

50 Logares R, Sunagawa S, Salazar G, Cornejo-Castillo FM, Ferrera I, Sarmento H, Hingamp P, Ogata H, de Vargas C, Lima-Mendez G, Raes J, Poulain J, Jaillon O, Wincker P, Kandels-Lewis S, Karsenti E, Bork P & Acinas SG (2014) Metagenomic 16S rDNA Illumina tags are a powerful alternative to amplicon sequencing to explore diversity and structure of microbial communities. Environmental Microbiology 16, 2659–2671.

51 de Vargas C, Audic S, Henry N, Decelle J, Mahe F, Logares R, Lara E, Berney C, Le Bescot N, Probert I, Carmichael M, Poulain J, Romac S, Colin S, Aury J-M, Bittner L, Chaffron S, Dunthorn M, Engelen S, Flegontova O, Guidi L, Horak A, Jaillon O, Lima-Mendez G, Luke J, Malviya S, Morard R, Mulot M, Scalco E, Siano R, Vincent F, Zingone A, Dimier C, Picheral M, Searson S, Kandels-Lewis S, Tara Oceans Coordinators, Acinas SG, Bork P, Bowler C, Gorsky G, Grimsley N, Hingamp P, Iudicone D, Not F, Ogata H, Pesant S, Raes J, Sieracki ME, Speich S, Stemmann L, Sunagawa S, Weissenbach J, Wincker P, Karsenti E, Boss E, Follows M, Karp-Boss L, Krzic U, Reynaud EG, Sardet C, Sullivan MB & Velayoudon D (2015) Eukaryotic plankton diversity in the sunlit ocean. Science 348, 1261605–1261605.

52. Martino C, Morton JT, Marotz CA, Thompson LR, Tripathi A, Knight R & Zengler K (2019) A Novel Sparse Compositional Technique Reveals Microbial Perturbations. mSystems 4, e00016-19.

53 Oksanen J (2018) Vegan: ecological diversity.

54 R Core Team (2010) R a language and environment for statistical computing: reference index R Foundation for Statistical Computing, Vienna.

55 Dinno A (2017) dunn.test: Dunn’s Test of Multiple Comparisons Using Rank Sums.

56 Tackmann J, Matias Rodrigues JF & von Mering C (2019) Rapid Inference of Direct Interactions in Large-Scale Ecological Networks from Heterogeneous Microbial Sequencing Data. Cell Systems 9, 286–296.e8.

57 Clauset A, Newman MEJ & Moore C (2004) Finding community structure in very large networks. Phys Rev E 70, 066111.

58 Frémont P, Gehlen M, Vrac M, Leconte J, Delmont TO, Wincker P, Iudicone D & Jaillon O (2022) Restructuring of plankton genomic biogeography in the surface ocean under climate change. Nat Clim Chang 12, 393–401.

59 Richter DJ, Watteaux R, Vannier T, Leconte J, Frémont P, Reygondeau G, Maillet N, Henry N, Benoit G, Da Silva O, Delmont TO, Fernàndez-Guerra A, Suweis S, Narci R, Berney C, Eveillard D, Gavory F, Guidi L, Labadie K, Mahieu E, Poulain J, Romac S, Roux S, Dimier C, Kandels S, Picheral M, Searson S, Tara Oceans Coordinators, Pesant S, Aury J-M, Brum JR, Lemaitre C, Pelletier E, Bork P, Sunagawa S, Lombard F, Karp-Boss L, Bowler C, Sullivan MB, Karsenti E, Mariadassou M, Probert I, Peterlongo P, Wincker P, de Vargas C, Ribera d’Alcalà M, Iudicone D & Jaillon O (2022) Genomic evidence for global ocean plankton biogeography shaped by large-scale current systems. eLife 11, e78129.

60 Brum JR, Ignacio-Espinoza JC, Roux S, Doulcier G, Acinas SG, Alberti A, Chaffron S, Cruaud C, de Vargas C, Gasol JM, Gorsky G, Gregory AC, Guidi L, Hingamp P, Iudicone D, Not F, Ogata H, Pesant S, Poulos BT, Schwenck SM, Speich S, Dimier C, Kandels-Lewis S, Picheral M, Searson S, Tara Oceans Coordinators, Bork P, Bowler C, Sunagawa S, Wincker P, Karsenti E & Sullivan MB (2015) Patterns and ecological drivers of ocean viral communities. Science 348, 1261498–1261498.

61 Ghiglione J-F, Galand PE, Pommier T, Pedrós-Alió C, Maas EW, Bakker K, Bertilson S, Kirchman DL, Lovejoy C, Yager PL & Murray AE (2012) Pole-to-pole biogeography of surface and deep marine bacterial communities. Proceedings of the National Academy of Sciences 109, 17633–17638.

62 De la Iglesia R, Echenique-Subiabre I, Rodríguez-Marconi S, Espinoza JP, von Dassow P, Ulloa O & Trefault N (2020) Distinct oxygen environments shape picoeukaryote assemblages thriving oxygen minimum zone waters off central Chile. Journal of Plankton Research 42, 514–529.

63 Schnetzer A, Moorthi SD, Countway PD, Gast RJ, Gilg IC & Caron DA (2011) Depth matters: Microbial eukaryote diversity and community structure in the eastern North Pacific revealed through environmental gene libraries. Deep Sea Research Part I: Oceanographic Research Papers 58, 16–26.

64 Parada V, Sintes E, van Aken HM, Weinbauer MG & Herndl GJ (2007) Viral Abundance, Decay, and Diversity in the Meso- and Bathypelagic Waters of the North Atlantic. Applied and Environmental Microbiology 73, 4429–4438.

65 Kaneko H, Blanc-Mathieu R, Endo H, Chaffron S, Delmont TO, Gaia M, Henry N, Hernández-Velázquez R, Nguyen CH, Mamitsuka H, Forterre P, Jaillon O, de Vargas C, Sullivan MB, Suttle CA, Guidi L & Ogata H (2021) Eukaryotic virus composition can predict the efficiency of carbon export in the global ocean. iScience 24, 102002.

66 Bettarel Y, Motegi C, Weinbauer MG & Mari X (2016) Colonization and release processes of viruses and prokaryotes on artificial marine macroaggregates. FEMS Microbiology Letters 363.

67 Durkin CA, Cetinić I, Estapa M, Ljubešić Z, Mucko M, Neeley A & Omand M (2022) Tracing the path of carbon export in the ocean though DNA sequencing of individual sinking particles. ISME J 16, 1896–1906.

68 Bates AE, Helmuth B, Burrows MT, Duncan MI, Garrabou J, Guy-Haim T, Lima F, Queiros AM, Seabra R, Marsh R, Belmaker J, Bensoussan N, Dong Y, Mazaris AD, Smale D, Wahl M & Rilov G (2018) Biologists ignore ocean weather at their peril. Nature 560, 299–301.

69 Mojica KDA & Brussaard CPD (2014) Factors affecting virus dynamics and microbial host–virus interactions in marine environments. FEMS Microbiology Ecology 89, 495–515.

70 Kukkaro P & Bamford DH (2009) Virus–host interactions in environments with a wide range of ionic strengths. Environmental Microbiology Reports 1, 71–77.

71 Bettarel Y, Bouvier T & Bouvy M (2009) Viral persistence in water as evaluated from a tropical/temperate cross-incubation. Journal of Plankton Research 31, 909–916.

72 Schulz F, Roux S, Paez-Espino D, Jungbluth S, Walsh DA, Denef VJ, McMahon KD, Konstantinidis KT, Eloe-Fadrosh EA, Kyrpides NC & Woyke T (2020) Giant virus diversity and host interactions through global metagenomics. Nature 578, 432–436.

73 Thamdrup B, Dalsgaard T & Revsbech NP (2012) Widespread functional anoxia in the oxygen minimum zone of the Eastern South Pacific. Deep Sea Research Part I: Oceanographic Research Papers 65, 36–45.

74 Berry D & Widder S (2014) Deciphering microbial interactions and detecting keystone species with co-occurrence networks. Front Microbiol 5.

75 Zhu W, Qin C, Ma H, Xi S, Zuo T, Pan W & Li C (2020) Response of protist community dynamics and co-occurrence patterns to the construction of artificial reefs: A case study in Daya Bay, China. Sci Total Environ 742, 140575.

76 Chaffron S, Delage E, Budinich M, Vintache D, Henry N, Nef C, Ardyna M, Zayed AA, Junger PC, Galand PE, Lovejoy C, Murray AE, Sarmento H, Tara Oceans coordinators, Acinas SG, Babin M, Iudicone D, Jaillon O, Karsenti E, Wincker P, Karp-Boss L, Sullivan MB, Bowler C, de Vargas C & Eveillard D (2021) Environmental vulnerability of the global ocean epipelagic plankton community interactome. Sci Adv 7, eabg1921.

77 Glock N, Roy A-S, Romero D, Wein T, Weissenbach J, Revsbech NP, Høgslund S, Clemens D, Sommer S & Dagan T (2019) Metabolic preference of nitrate over oxygen as an electron acceptor in foraminifera from the Peruvian oxygen minimum zone. PNAS 116, 2860–2865.

## References

Adriaenssens, EM, and Cowan, DA. Using signature genes as tools to assess environmental viral ecology and diversity. AEM, AEM-00878, 2013.

Bateman, A et al. “The Pfam protein families database.” Nucleic acids research 32.suppl_1 D138-D141, 2004:.

Brum JR, et al. Patterns and ecological drivers of ocean viral communities. Science 348: 1261498–1261498, 2015.

Callahan, BJ, McMurdie, PJ, and Holmes, SP. Exact sequence variants should replace operational taxonomic units in marker-gene data analysis. The ISME journal 2639–2643, 2017.

Clarke, KR. Non-parametric multivariate analyses of changes in community structure. Aust J Ecol 18:117–43, 1993.

de Vargas C, et al. Eukaryotic plankton diversity in the sunlit ocean. Science 348: 1261605– 1261605, 2015.

Del Campo, Javier, et al. EukRef: phylogenetic curation of ribosomal RNA to enhance understanding of eukaryotic diversity and distribution. PLoS biology 16.9 e2005849, 2018.

Edgar, Robert. *Usearch*. Lawrence Berkeley National Lab. (LBNL), Berkeley, CA (United States), 2010.

Endo H, Blanc-Mathieu R, Li Y, Salazar G, Henry N, Labadie K, et al. Biogeography of marine giant Phageses reveals their interplay with eukaryotes and ecological functions. Nat Ecol Evol 4: 1639–1649, 2020.

Enright, AJ, Van Dongen, S, Ouzounis, CA. An efficient algorithm for large-scale detection of protein families. Nucleic acids research 30.7 1575–1584, 2002.

Gregory AC, Zayed AA, et al. Marine DNA Viral Macro- and Microdiversity from Pole to Pole. Cell 177: 1109–1123.e14, 2019.

Guillou, L, et al. The Protist Ribosomal Reference database (PR2): a catalog of unicellular eukaryote small sub-unit rRNA sequences with curated taxonomy. Nucleic acids research 41.D1 D597–D604, 2012.

Hyatt, D, et al. Prodigal: prokaryotic gene recognition and translation initiation site identification. BMC bioinformatics 11.1, 119, 2010.

Ibarbalz FM, et al. Global Trends in Marine Plankton Diversity across Kingdoms of Life. Cell 179: 1084–1097.e21, 2019.

Kanehisa, M, et al. The KEGG databases at GenomeNet. Nucleic acids research 30.1, 42–46, 2002.

Katoh, K, and Standley, DM. MAFFT multiple sequence alignment software version 7: improvements in performance and usability. Mol. Biol. Evol. 30, 772–780, 2013.

Kurtz, S, et al. Versatile and open software for comparing large genomes. Genome biology 5.2, 1–9, 2004.

Li, W., and Godzik, A. Cd-hit: a fast program for clustering and comparing large sets of protein or nucleotide sequences. Bioinformatics 22.13, 1658–1659, 2006.

Logares R, et al. Metagenomic 16S rDNA Illumina tags are a powerful alternative to amplicon sequencing to explore diversity and structure of microbial communities. Environ Microbiol. Sep;16(9):2659–71, 2014.

Mahé, F, et al. Swarm v2: highly-scalable and high-resolution amplicon clustering. PeerJ 3 e1420, 2015.

Matsen, FA, Kodner, RB, Armbrust, EV. pplacer: linear time maximum-likelihood and Bayesian phylogenetic placement of sequences onto a fixed reference tree. BMC Bioinform. 11, 538, 2010.

Quast, C., et al. The SILVA ribosomal RNA gene database project: improved data processing and web-based tools. Nucleic Acids Res 41: D590–D596, 2013.

Stamatakis, A. RAxML-VI-HPC: maximum likelihood-based phylogenetic analyses with thousands of taxa and mixed models. Bioinformatics 22, 2688–2690, 2006.

Suzek, BE, et al. UniRef: comprehensive and non-redundant UniProt reference clusters. Bioinformatics 23.10, 1282–1288, 2007.

Zdobnov, EM, and Apweiler, R. InterProScan–an integration platform for the signature-recognition methods in InterPro. Bioinformatics 17.9, 847–848, 2001.

